# Ketolysis is a metabolic driver of CD8^+^ T cell effector function through histone acetylation

**DOI:** 10.1101/2022.08.26.505402

**Authors:** Katarzyna M. Luda, Susan M. Kitchen-Goosen, Eric H. Ma, McLane J. Watson, Lauren R. Duimstra, Brandon M. Oswald, Joseph Longo, Zhen Fu, Zachary Madaj, Ariana Kupai, Bradley M. Dickson, Irem Kaymak, Kin H. Lau, Shelby Compton, Lisa M. DeCamp, Daniel P. Kelly, Patrycja Puchalska, Kelsey S. Williams, Connie M. Krawczyk, Dominique Lévesque, François-Michel Boisvert, Ryan D. Sheldon, Scott B. Rothbart, Peter A. Crawford, Russell G. Jones

## Abstract

Environmental nutrient availability influences T cell metabolism, impacting T cell function and shaping immune outcomes. However, the metabolic pathways critical for optimal T cell responses remain poorly understood. Here, we identify ketone bodies (KBs) – including β-hydroxybutyrate (βOHB) and acetoacetate (AcAc) – as essential fuels supporting CD8^+^ T cell metabolism and effector function. Ketolysis is an intrinsic feature of highly functional CD8^+^ T effector (Teff) cells and βOHB directly increases CD8^+^ Teff cell IFN-γ production and cytolytic activity. Using metabolic tracers, we establish that CD8^+^ Teff cells preferentially use KBs over glucose to fuel the tricarboxylic acid (TCA) cycle *in vitro* and *in vivo*. KBs directly boost the respiratory capacity of CD8^+^ T cells and TCA cycle-dependent metabolic pathways that fuel T cell growth. Mechanistically, we find that βOHB is a major substrate for acetyl-CoA production in CD8^+^ T cells and regulates effector responses through effects on histone acetylation. Together, our results identify cell-intrinsic ketolysis as a metabolic and epigenetic driver of optimal CD8^+^ T cell effector responses.

**One Sentence summary:** Ketone bodies promote CD8^+^ T cell metabolism and effector function through regulation of epigenetic programming

## Main text

Multi-cellular organisms have evolved protective strategies for defending against pathogens such as viruses and bacteria and intrinsic threats such as cancer. Central to host defense are CD8^+^ T cells, which are critical for pathogen clearance and protecting the host from re-infection through long-lived immune memory (*1, 2*). One of the fundamental biological programs supporting T cell effector function is cellular metabolism, which generates energy and biosynthetic precursors essential for CD8^+^ T effector (Teff) cell proliferation, survival, and production of effector molecules (i.e., IFN-γ, TNF-α, and cytolytic factors) (*3–5*). CD8^+^ Teff cells undergo extensive metabolic rewiring to arm themselves with the bioenergetic and biosynthetic capacity to successfully protect the host (*6, 7*). However, Teff cell function is highly dependent on nutrient availability (*8–11*), which in the whole organism is governed by diet and host metabolism. Changes in host metabolic homeostasis triggered by infection−which can include catabolic wasting (“cachexia”) and anorexia−can influence the pathogenesis of the infection and affect disease tolerance (*12, 13*). In response to bacterial and viral infections, disrupted feeding behavior and metabolic rewiring in tissues, such as the liver, promote changes in host metabolism including lipolysis and production of ketone bodies (*14–17*); however, how T cell-intrinsic metabolic programming synergizes with changes in host metabolism during an immune response remains poorly defined. Here, we set out to delineate cell-intrinsic and host metabolic factors critical for the formation of highly functional CD8^+^ Teff cells.

Effective control of infection and cancer growth requires robust effector CD8^+^ T cell responses, which are driven by TCR-dependent transcriptional, epigenetic, and metabolic programs (*2, 18–20*). To define features of highly functional CD8^+^ T effector (Teff) cells from setting of both cancer and infection, we conducted a meta-analysis of three independent studies characterizing gene expression profiles of CD8^+^ T cells responding to acute infection (*Listeria monocytogenes* (*Lm*) or *Lymphocytic choriomeningitis virus* (LCMV), Armstrong strain), chronic infection (LCMV, Clone 13 (CL-13) strain), or syngeneic tumors (*21–23*). Pearson correlations (**Fig. 1a, S1a**) and principal component analysis (PCA) among samples revealed clustering of samples into 6 main cellular subtypes: 1) naïve (Tn), 2) Teff, 3) T memory (Tmem), 4) early cancer T exhausted (Tex), 5) late cancer Tex, and 6) chronic virus Tex. To identify genes preferentially associated with “functional” versus “dysfunctional” CD8^+^ T cells, we compared gene expression patterns between Teff cells and late cancer- or chronic viral-driven Tex cells (**Table S1**). Wald test statistics for each gene were generated for each pairwise comparison (cancer and viral datasets) and median scores used to identify genes associated with functional versus dysfunctional states (**Fig. 1b**). Rank analysis revealed that genes associated with CD8^+^ T cell cytotoxicity (i.e., *Klrg1, Klf2, Rora, Lef1*) were enriched in the functional Teff cell cohort, while dysfunctional CD8^+^ T cells displayed enrichment in genes associated with T cell exhaustion (i.e., *Tox, Pdcd1, Entpd1, Lag3*) (**Fig. 1b, Fig. S1b, Table S1**) and inflammatory signaling (**Fig. S1c**). Given the central role of metabolism in supporting Teff cell responses (*6, 24*), we hypothesized that specific metabolic pathways may contribute to CD8^+^ Teff cell function. KEGG pathway analysis of the top ∼4% of genes expressed by functional CD8^+^ Teff cells revealed preferential enrichment of several metabolic pathways (**Fig. 1c**). Among these, the synthesis and degradation of ketone bodies was the most highly enriched metabolic pathway in functional CD8^+^ T cells (**Fig. 1c**).

**Figure 1.**
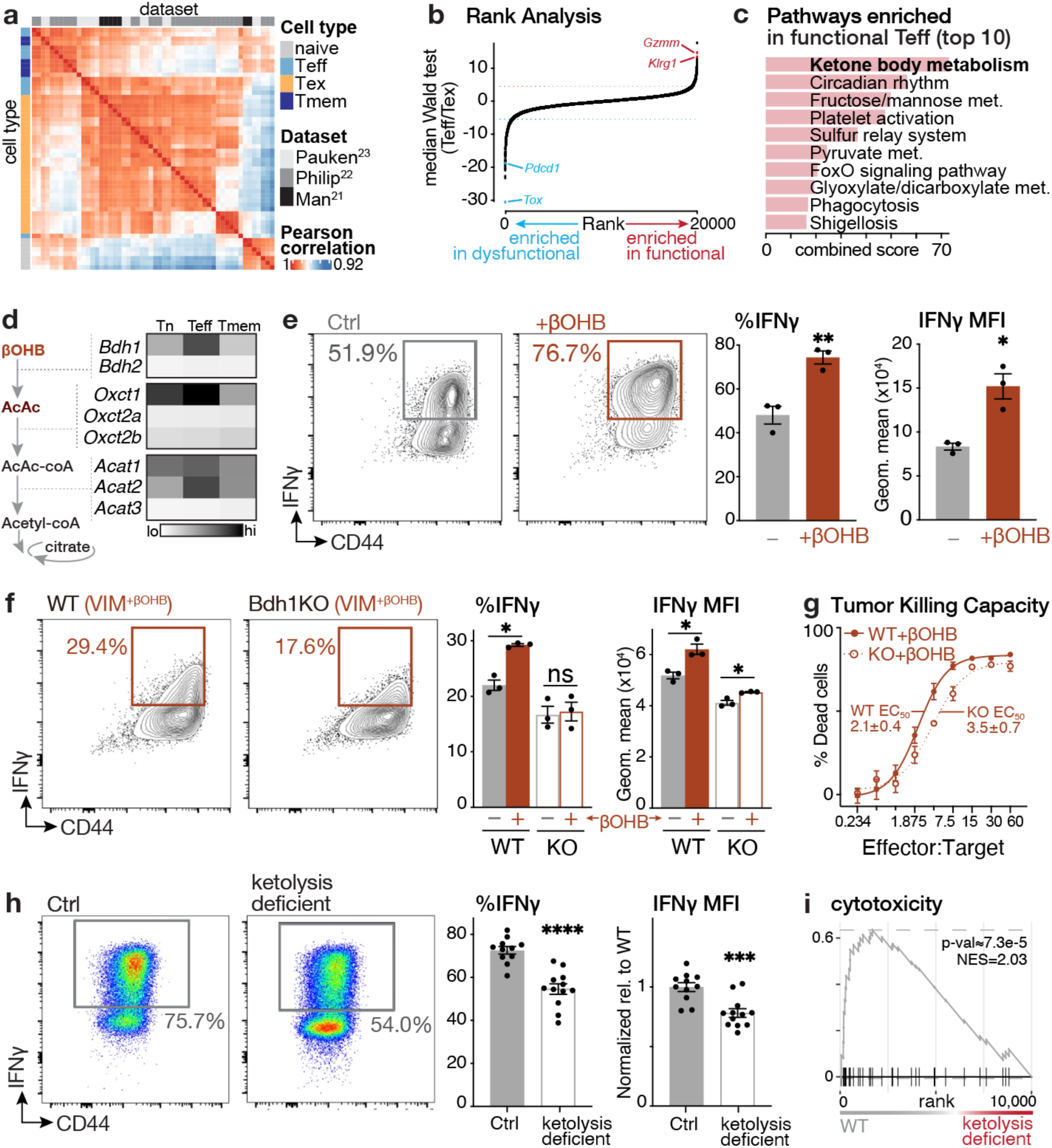
Ketolysis is a metabolic feature of functional CD8^+^ effector T cells. (**a**) Pearson correlation-driven similarity matrix analysis of gene expression profiles of CD8^+^ T cell states (naïve, Teff – effector T cells, Tex – exhausted T cells, Tmem – memory T cells). Analysis was conducted on RNA-seq datasets from three independent studies characterizing gene expression profiles of antigen-specific CD8^+^ T cells from acute infection (LCMV Armstrong, *Lm*), chronic infection (LCMV CL-13), and cancer (hepatocellular carcinoma) models *in vivo*. (**b**) Rank analysis of genes enriched in dysfunctional and in functional CD8^+^ T cell states. Median Wald test statistics for DEG between Teff and Tex populations (Teff/Tex) were calculated based on cancer and virus response datasets. (**c**) Pathway analysis of the top 10 KEGG pathways enriched in functional Teff cells from (b). (**d**) Heatmap of ketolytic gene expression in OT-I CD8^+^ naïve (Tn) or OT-I CD8^+^ Teff and Tmem Ocells triggered in response to *Lm*-OVA infection (Teff, 2 dpi; Tmem, 30 dpi). A schematic of the enzymes involved in ketolysis is shown. (**e**) IFN-γ production by CD8^+^ T cells cultured in IMDM in the presence (+βOHB) or absence (Ctrl) of 5 mM βOHB. *Left,* Representative flow cytometry plots for CD44 versus IFN-γ expression by CD8^+^ T cells 3 days post activation. *Right,* Bar graph showing the percentage of IFN-γ^+^ T cells and the geometric mean fluorescence intensity (MFI) for IFN-γ^+^ T cells cultured with (+) or without (-) βOHB. Data represent the mean ± SEM (n=3). (**f**) IFN-γ production by wild type (WT) and Bdh1-deficient (Bdh1KO) CD8^+^ T cells cultured in VIM containing or lacking 5 mM βOHB. *Left,* Representative flow cytometry plots for CD44 versus IFN-γ expression by WT and Bdh1KO CD8^+^ T cells cultured with βOHB 3 days post activation. *Right,* Bar graph showing the percentage of IFN-γ ^+^ T cells and the geometric mean fluorescence intensity (MFI) for IFN-γ ^+^ T cells cultured with (+) or without (-) βOHB. Data represent the mean ± SEM (n=3). (**g**) *In vitro* killing assay for WT and Bdh1KO T cells cultured in the presence of βOHB. Shown is the percentage of dead MC38-OVA tumor cells after 48 h of co-culture with activated WT or Bdh1KO OT-I CD8^+^ cells at the indicated effector:target (E:T) ratio. The E:T ratio required to kill 50% tumor cells (EC_50_) for each genotype is indicated (mean ± SEM, n=3). (**h**) IFN-γ production by control (Ctrl) and ketolysis-deficient OT-I CD8^+^ cells isolated from *Lm*-OVA-infected mice (7 dpi). *Left,* Representative CD44 versus IFN-γ expression for control and ketolysis-deficient OT-I cells. *Right,* Percentage of IFN-γ^+^ OT-I T cells and MFI of IFN-γ expression (normalized to Ctrl cells) for OT-I cells isolated from the spleen of *Lm*-OVA-infected mice 7 dpi (mean±SEM, n=11-12/group). (**i**) Gene set enrichment analysis (GSEA) of CD8^+^ T cell cytotoxicity genes in control versus ketolysis-deficient OT-I T cells responding to *Lm*-OVA (7 dpi, n=3 biological replicates/group). Statistical significance and normalized enrichment scores (NES) are indicated. *, *p* < 0.05; **, *p* < 0.01; ***, *p* < 0.001; ****, p < 0.0001; ns, not significant.

Ketone bodies (KBs), including β-hydroxybutyrate (βOHB) and acetoacetate (AcAc), are alternative metabolic fuels critical for supporting bioenergetic metabolism during periods of nutrient deprivation (*25–27*). KBs can also have fates beyond terminal oxidation, including impact on cell signaling, direct and indirect effects on histone post-translational modifications (PTMs), and inhibition of inflammatory processes (*26, 28*). The primary site of KB production in mammals is the liver, which generates AcAc and βOHB from fatty acid-derived acetyl-CoA. KBs are subsequentially transported to extrahepatic tissues for terminal oxidation in the tricarboxylic acid (TCA) cycle. KBs are metabolized to acetyl-CoA in a set of enzymatic reactions collectively called ketolysis. Mitochondrial βOHB dehydrogenase (BDH1) converts D-βOHB to AcAc, which is in turn converted to acetoacetyl-CoA (AcAc-CoA) by succinyl-CoA-3-oxaloacid CoA transferase (SCOT, encoded by the *Oxct* genes) (**Fig. 1d**). AcAc-CoA is further processed by thiolase acetyl-CoA acetyltransferase (ACAT) to generate two molecules of acetyl-CoA, which can enter the TCA cycle (**Fig. 1d**). Gene expression analysis of CD8^+^ T cells responding to *Lm* infection (*29*) revealed increased expression of mRNA transcripts encoding several enzymes in the pathway (i.e., *Bdh1, Oxct1, Acat1/2*) in Teff cells relative to Tn and Tmem cells (**Fig. 1d**). Proteomic analysis of CD8^+^ T cells (*30, 31*) revealed a ∼2-3-fold induction in BDH1 and SCOT in Teff cells following *in vitro* activation (**Fig. S2a**), while Teff cells responding to *Lm* infection *in vivo* displayed prominent increases in BDH1 expression (**Fig. S2b**). Thus, the transition from naïve to effector T cell states is associated with increased expression of ketolytic enzymes.

We next asked if KBs could exert direct impact on CD8^+^ Teff cell function. CD8^+^ Teff cells activated in the presence of D-βOHB (5 mM) displayed increased capacity to produce IFN-γ, reflected by both increased numbers of IFN-γ-producing cells and increased IFN-γ protein expression per cell (as determined by increased mean fluorescence intensity (MFI) of IFN-γ staining) (**Fig. 1e**). We recently reported that the presence of physiologic carbon sources (PCS) enhances CD8^+^ T cell effector function, including IFN-γ and granzyme production (*32*). Removing βOHB from PCS-supplemented cell culture medium reduced IFN-γ production by CD8^+^ T cells, while addition of βOHB alone was sufficient to increase IFN-γ levels, suggesting that βOHB was the active metabolite in PCS acting to enhance IFN-γ production (**Fig. S3a**). Next, we assessed the function of CD8^+^ T cells lacking BDH1 expression (via crossing of *Bdh1*-floxed mice (*33*) to *Cd4-Cre* transgenic mice to generate conditional deletion of BDH1 in mature T cells). BDH1-deficient (Bdh1KO) CD8^+^ Teff cells displayed reduced IFN-γ production relative to control cells following *in vitro* activation (**Fig. S3b**), and BDH1 was required for the βOHB-driven increase in IFN-γ production by CD8^+^ Teff cells (**Fig. 1f**). In addition, Bdh1KO T cells displayed a 30-40% reduction in their ability to lyse MC38 tumor cells (**Fig. 1g**), implicating ketolysis as a regulator of cytolytic capacity.

Finally, we assessed the role of cell-intrinsic ketolysis in CD8^+^ Teff cell function *in vivo* following infection with *Lm*-OVA, which induces robust expansion of IFN-γ-producing CD8^+^ T cells (*34*). Given that both βOHB and AcAc are present in circulation (*35*) and that either metabolite may mediate the effects of KBs on T cell function, we generated ketolysis-deficient T cells via shRNA-mediated silencing *of Oxct1* in Bdh1KO T cells (validation in **Fig. S4a**). OVA-specific OT-I^+^ CD8^+^ T cells (Control or Bdh1KO) were transduced with control (*shFF*, targeting firefly luciferase) or *Oxct1*-targeting retroviral vectors, transferred into congenic hosts, followed by infection with *Lm*-OVA. CD8^+^ Teff cell responses were then analyzed 7 days post infection (dpi) (**Fig. S4b-c**). Ketolysis-deficient OT-I T cells displayed increased expansion *in vivo* (**Fig. S4d**) but no major change in effector or memory precursor subsets (**Fig. S4e**). However, similar to our observation with *in vitro*-activated T cells (**Fig. 1f, S3b**), ketolysis-deficient CD8^+^ T cells displayed lower IFN-γ production in vivo compared to control cells (**Fig. 1h**). To identify the molecular mechanisms by which ketolysis drives CD8^+^ T cell effector function, we performed transcriptomics on control and ketolysis-deficient OT-I T cells isolated from the spleens of *Lm*-OVA-infected mice 7 dpi (**Table S2, Fig. S5a**). Pathway analysis revealed enrichment of proliferative programs (i.e., cell cycle, E2F targets)−consistent with increased expansion of ketolysis-deficient OT-I cells *in vivo* (**Fig. S4d**)−and a loss of inflammatory signatures (i.e., IFN-α/IFN-γ response, allograft rejection) in ketolysis-deficient T cells (**Fig. S5b-c**). Strikingly, gene set enrichment analysis (GSEA) using gene signatures derived from single cell profiling of human T cells (**Table S3**) (*36*) revealed a loss of the CD8^+^ cytotoxicity signature in ketolysis-deficient T cells (**Fig. 1i**). Collectively, these findings suggest that cell-intrinsic ketolysis is required for the development of CD8^+^ T cell functional programs.

Given that KBs play critical roles in organismal energy homeostasis, including serving as oxidative fuels during states of low nutrient availability (*26*), we next questioned whether KBs could function as a fuel source for CD8^+^ T cells. Using ^13^C-based metabolic tracers, we found that D-[U-^13^C_4_]βOHB was readily imported by proliferating CD8^+^ Teff cells, saturating the intracellular βOHB pool within 2 minutes of exposure (**Fig. 2a, S6a**). While previous work suggests that Tmem cells can undergo ketogenesis (*37*), rapid and near complete ^13^C-enrichment of the βOHB pool from [U-^13^C_4_]βOHB suggests the absence of ketogenesis from endogenous substrates in Teff cells, which would dilute the total βOHB pool with unlabeled βOHB. Furthermore, as expected, we did not observe contribution of [U-^13^C_6_]glucose to intracellular βOHB in Teff cells (**Fig. 2a**). CD8^+^ T cells display plasticity in metabolic fuel choice, which allows them to operate and maintain effector function in diverse metabolic environments (*5*). To compare the contribution of KB-derived carbon with other carbon sources for TCA cycle metabolism, we cultured CD8^+^ Teff cells in the presence of ^13^C-labelled substrates at concentrations commonly observed in plasma (*38, 39*) and assessed contribution to TCA cycle-derived metabolites. Of all ^13^C-labeled substrates, [U-^13^C_4_]βOHB-derived carbon contributed the highest enrichment to TCA cycle intermediates (**Fig. 2b**), contributing to the citrate (M+2) pool 5-fold more effectively than [U-^13^C_6_]glucose (**Fig. S6b**), indicating that βOHB could contribute to TCA cycle metabolism even under nutrient-replete conditions.

**Figure 2.**
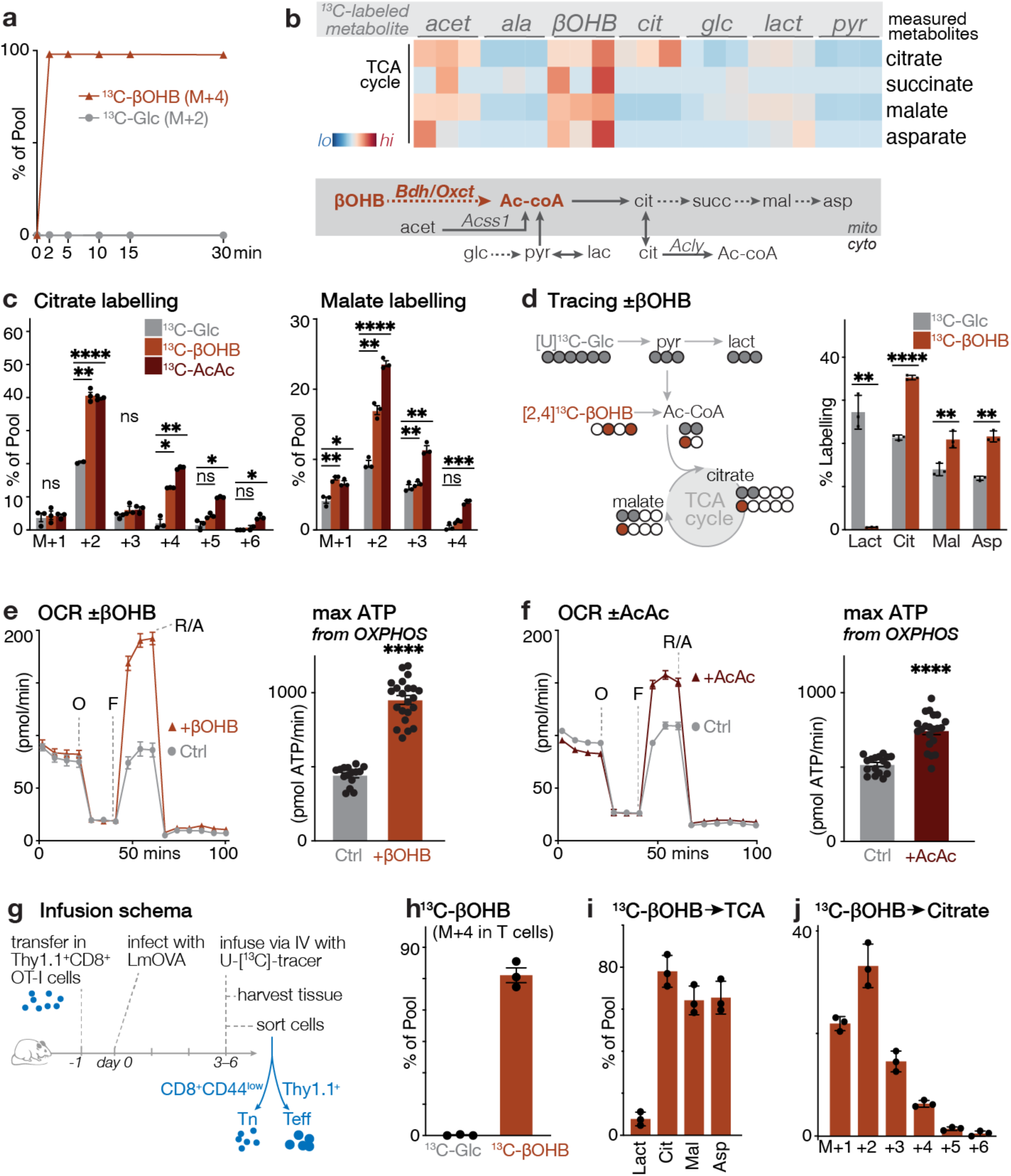
Ketone bodies are physiologic fuels for CD8^+^ effector T cells. (**a**) Timecourse of βOHB uptake and cell-intrinsic βOHB production from glucose in activated CD8^+^ T cells. Shown is the contribution of ^1^[^13^C_4_]βOHB (M+4, from exogenous [^13^C_4_]βOHB) or [U-^13^C_6_]glucose-derived βOHB (M+2) to the overall intracellular βOHB pool over time (mean ± SEM, n=3). (**b**) Heatmap representing relative contribution of ^13^C from indicated ^13^C-labelled substrates into TCA cycle-derived metabolites. Shown is a schematic depicting the contribution of different carbon sources to TCA cycle metabolism, with enzymes and enzyme reactions localized to the cytosol and mitochondrion indicated. (**c**) Mass isotopologue distribution (MID) for [U-^13^C_6_]glucose-, [^13^C_4_]βOHB-, and [^13^C_4_]AcAc-derived carbon into citrate and malate for activated CD8^+^ Teff cells after 2 h of culture (n=3/group). (**d**) Direct comparison of TCA cycle labeling from βOHB and glucose using competitive tracing. Activated CD8^+^ Teff cells were cultured in medium containing 5 mM [U-^13^C_6_]glucose and 2 mM [2,4-^13^C_2_]βOHB for 2 h and the relative contribution of ^13^C-label from [U-^13^C_6_]glucose (M+2) or [2,4-^13^C_2_]βOHB (M+1) to TCA cycle metabolite pools is shown (mean ± SEM, n=3). Lactate labeling from [U-^13^C_6_]glucose (M+3) or [2,4-^13^C_4_]βOHB (M+1) is shown as a control. (**e-f**) Bioenergetic profile of *in vitro*-activated CD8^+^ T cells cultured with **(e)** 2 mM βOHB or **(f)** 2 mM AcAc (mean ± SEM, n=20–24/group). *Left*, OCR plots for activated CD8^+^ T cells over time for βOHB and AcAc. Time of addition of oligomycin (O), fluoro-carbonyl cyanide phenylhydrazone (FCCP, F), and rotenone and antimycin A (R/A) are indicated. *Right,* maximal ATP production rates from OXPHOS following addition of βOHB and AcAc. T cells that received no exogenous substrates (Ctrl) are indicated. (**g**) Schematic of experimental set up for ^13^C infusions in *Lm*-OVA-infected mice using [^13^C_4_]βOHB and [U-^13^C_6_]glucose. (**h**) Relative contribution of infused [U-^13^C_6_]glucose or [^13^C_4_]βOHB to the intracellular βOHB (M+4) pool in *Lm*-OVA-specific OT-I T cells 6 dpi (mean ± SEM, n = 3). ^13^C metabolite enrichment was normalized relative to [U-^13^C_6_]glucose (M+6) or [^13^C_4_]βOHB (M+4) levels in serum. (**i-j**) Enrichment of infused [^13^C_4_]βOHB-derived carbon into intracellular metabolites in *Lm*-OVA-specific OT-I T cells 6 dpi (mean ± SEM, n = 3). **(j)** Total ^13^C enrichment from [^13^C_4_]βOHB in lactate, citrate, malate, and aspartate. **(j)** MID of [^13^C_4_]βOHB-derived carbon in intracellular citrate. ^13^C metabolite enrichment was normalized relative to [^13^C_4_]βOHB (M+4) levels in serum. *, *p* < 0.05; **, *p* < 0.01; ***, *p* < 0.001; ****, p < 0.0001; ns, not significant.

We continued our *in vitro* studies using KBs at 2 mM (unless stated otherwise) to mimic the concentration of circulating βOHB achieved with therapeutic ketogenic diets (*40, 41*). Both [U-^13^C_4_]βOHB and [U-^13^C_4_]AcAc significantly contributed to citrate and malate synthesis even in the presence of physiologic glucose levels (5 mM) (**Fig. 2c**). Of note, the ratio of malate (M+2) to citrate (M+2) labeling from ^13^C-βOHB was approximately 0.5, indicative of “non-canonical” TCA cycle activity (i.e., the citrate-malate shuttle) (*42*) and potential use of citrate for other reactions beyond the TCA cycle (i.e., export to the cytosol for acetyl-CoA production). To assess direct competition between glucose and βOHB for TCA cycle metabolism, we designed a co-labeling experiment where activated CD8^+^ Teff cells were cultured with [U-^13^C_6_]glucose and [2,4-^13^C_2_]βOHB. In this setting, metabolic intermediates generated during the first turn of the TCA cycle would be labelled M+1 from βOHB and M+2 from glucose (**Fig. 2d**). As expected, ^13^C-labeled lactate (M+3) was generated exclusively from [U-^13^C_6_]glucose and not [2,4-^13^C_2_]βOHB. (**Fig. 2d**); however, βOHB contributed approximately 50% more carbon to the synthesis of TCA cycle intermediates−particularly citrate−compared to glucose, despite being at lower molar concentration (**Fig. 2d**). These observations are highly analogous to macrophages that preferentially use AcAc over glucose for TCA cycle metabolism (*35*). Collectively, these data confirm that KBs are metabolic substrates for CD8^+^ T cells, and that βOHB is preferentially used by T cells over glucose to supply TCA cycle metabolism, even under nutrient-replete conditions in which glucose is abundant.

Given the contribution of βOHB and AcAc to TCA cycle metabolism, we next examined the impact of KB metabolism on T cell bioenergetics. Short-term exposure (2h) of activated CD8^+^ Teff cells to either βOHB (**Fig. 2e**) or AcAc (**Fig. 2f**) modestly increased their basal oxygen consumption rate (OCR) and ATP production from OXPHOS (**Fig. S6c**). However, the maximal respiratory rate of CD8^+^ T cells was greatly increased by KBs (2-fold and 1.5-fold increases for βOHB and AcAc, respectively), which corresponded to a significant increase in their maximal ATP production rate from OXPHOS (**Fig. 2e-f**). Collectively, these data indicate that KBs directly augment mitochondrial ATP production by boosting maximal respiratory capacity.

Finally, we used *in vivo* ^13^C metabolite infusions to evaluate whether T cells use KBs as a fuel source *in vivo*. We transferred Thy1.1^+^ OT-I CD8^+^ T cells into congenic recipient mice, infected the mice with *Lm*-OVA one day later, and then infused the mice for 2 h at 2 dpi (during proliferative expansion) or 6 dpi (peak of T cell response) with D-[^13^C_4_]βOHB or [U-^13^C_6_]glucose prior to CD8^+^ T cell isolation and metabolomic analysis (**Fig. 2g**) (*43, 44*). With our [U-^13^C_4_]βOHB infusion strategy, we achieved ∼75% enrichment of fully labeled (M+4) ^13^C-labeled βOHB in circulation at both 2 and 6 dpi (**Fig. S7a-b**), with βOHB plasma levels achieving similar levels as observed in ketogenic diet settings (∼2 mM). Moreover, we detected M+4 ^13^C-βOHB in *Lm*-OVA-specific CD8^+^ T cells after only 2 h of infusion (**Fig. S7b**), corresponding to ∼80-85% labelling of the total βOHB pool (**Fig. 2h**), indicating rapid import of βOHB by CD8^+^ Teff cells *in vivo*. While we achieved good enrichment (∼40%) of fully labeled (M+6) ^13^C-glucose in the circulating glucose pool (**Fig. S7c**), we observed no contribution of glucose to βOHB production in T cells (**Fig. 2h**) or circulating βOHB levels in plasma (**Fig. S7d**). Similar to our observations in vitro (**Fig. 2d**), infused [U-^13^C_4_]βOHB readily labeled TCA cycle and TCA cycle-derived metabolites in *Lm*-OVA-specific CD8^+^ T cells *in vivo* (**Fig. 2i**).

Higher-order labeling patterns for [U-^13^C_4_]βOHB in citrate (i.e., M+3, M+4) suggests that βOHB carbon contributed to several turns of the TCA cycle within the short 2 h infusion period (**Fig. 2j**). Finally, we observed several key trends when we compared *in vivo* utilization of [U-^13^C]βOHB to [U-^13^C_6_]glucose by CD8^+^ T cells. When normalized to circulating infusion rates, both βOHB and glucose displayed similar contribution of ^13^C-carbon to the TCA cycle (M+2 citrate, malate and aspartate) during early T cell expansion *in vivo* (2 dpi); however, while glucose utilization by T cells declined by the peak of the T cell response to *Lm*-OVA (6 dpi), [^13^C_4_]βOHB still readily labeled intermediates of the TCA cycle (**Fig. S7e-f**), suggesting preferred use of βOHB as a TCA cycle fuel during the peak effector phase of the T cell response. Collectively, these data indicate that βOHB is utilized as a preferred fuel by CD8^+^ Teff cells both *in vitro* and *in vivo* during an active immune response, even when glucose is present.

Given our results that ketolysis can fuel CD8^+^ T cell metabolism, we hypothesized that the contribution of ketones to mitochondrial metabolism and acetyl-CoA production may underlie the effects of KBs on T cell function. The bioenergetic potential from KBs comes from both the dehydrogenation of βOHB to AcAc by BDH1, which directly generates NADH, and the production of acetyl-CoA that can enter the TCA cycle and fuel downstream ATP production by the electron transport chain (**Fig. 3a**). Consistent with this, we found that BDH1 was required for processing of [U-^13^C_4_]βOHB but not [U-^13^C_4_]AcAc into citrate (**Fig. 3b**). As expected, targeting the downstream ketolytic enzyme SCOT (via silencing of *Oxct1*) strongly diminished production of citrate from both [U-^13^C_4_]βOHB and [U-^13^C_4_]AcAc (**Fig. 3c**). Surprisingly, using a Seahorse Bioanalyzer, we observed that both the increased respiratory capacity (**Fig. 3d**) and βOHB-induced boost in both basal and maximal ATP production from OXPHOS (**Figs. 3e, S8**) was dependent on BDH1 but not on SCOT: βOHB was still able to increase ATP production from OXPHOS in SCOT-deficient T cells (**Fig. 3e**). Collectively, these data indicate that in CD8^+^ Teff cells BDH1 is a critical regulator of both NADH production for OXPHOS and citrate synthesis for downstream metabolic reactions.

**Figure 3.**
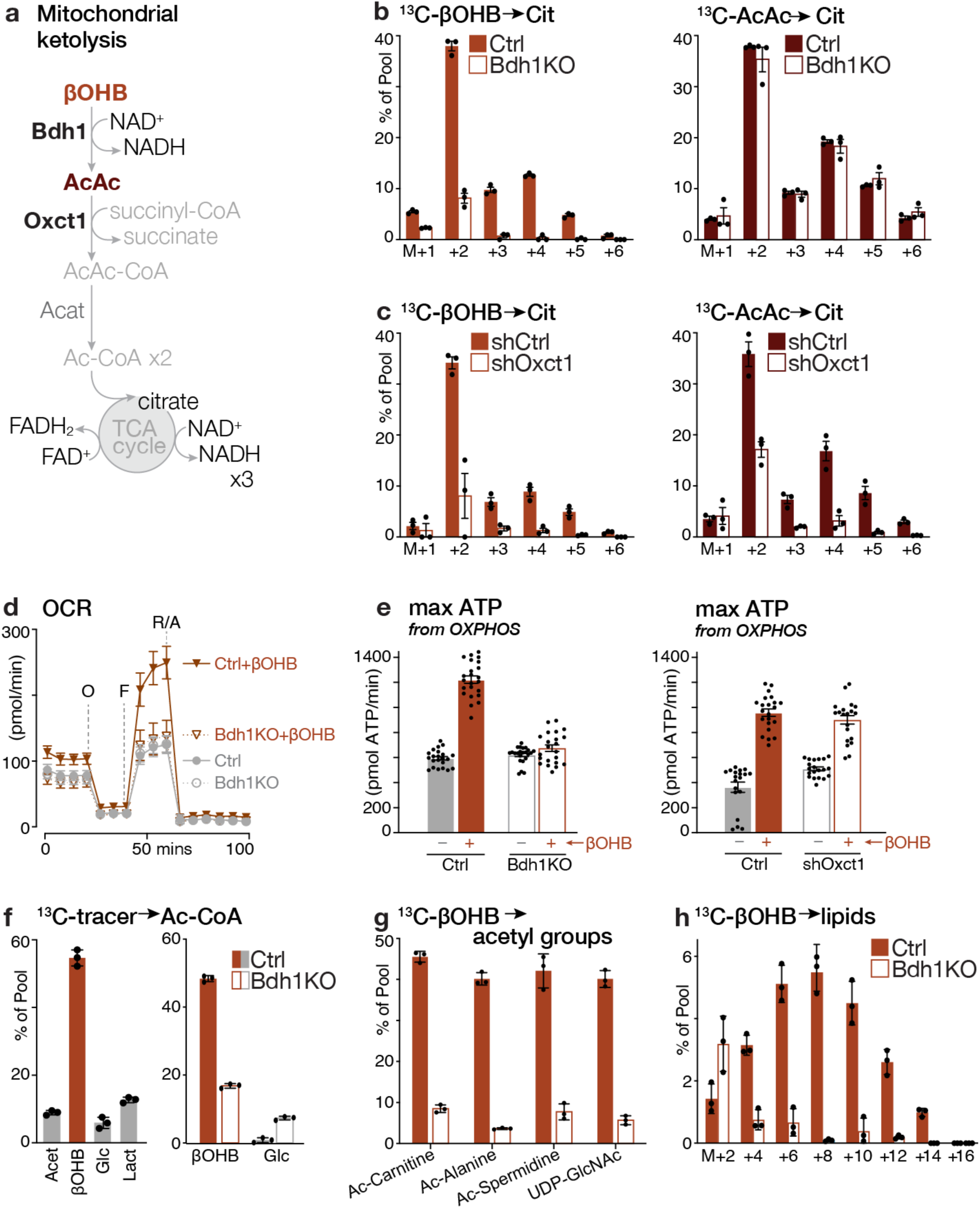
Cell-intrinsic ketolysis regulates CD8^+^ effector T cell bioenergetics and acetyl-CoA production. (**a**) Illustration of the mitochondrial ketolysis pathway. (**b**) Labeling of 2 mM [^13^C_4_]βOHB (left) or [^13^C_4_]AcAc (right) into intracellular citrate in activated control (Ctrl) or Bdh1-deficient (Bdh1KO) CD8^+^ T cells after 2 h of culture (mean ± SEM, n=3/group). (**c**) [^13^C_4_]βOHB (left) or [^13^C_4_]AcAc (*right*) labeling into intracellular citrate in CD8^+^ T cells expressing control (*shCtrl*) or Oxct1-targeting (*shOxct1*) shRNAs (mean ± SEM, n=3/group). Cells were cultured with ^13^C substrates for 2 h as in (b). (**d-e**) Bioenergetic profile of ketolysis deficient CD8^+^ T cells. **(d)** OCR plot over time for activated control (Ctrl) or Bdh1KO CD8^+^ T cells cultured for 2h without or with 2 mM βOHB. Time of addition of oligomycin (O), FCCP (F), and rotenone and antimycin A (R/A) are indicated. **(e)** Maximal ATP production rates from OXPHOS in activated Bdh1KO (left) or *shOxct1*-expressing (right) CD8^+^ T cells and respective controls (Ctrl) following addition of 2 mM βOHB. (**f**) Metabolic production of acetyl-CoA in CD8^+^ T cells. *Left,* Fractional enrichment of [^13^C_2_]acetate, [^13^C_4_]βOHB, [^13^C_6_]glucose, and [^13^C_3_]lactate in the intracellular acetyl-CoA (M+2) pool of activated CD8^+^ T cells following 24 h of culture (mean ± SEM, n=3). *Right,* Fractional enrichment of [^13^C_4_]βOHB and [^13^C_6_]glucose carbon into the acetyl-CoA (M+2) pool in control (Ctrl) and Bdh1KO CD8^+^ T cells after 24 h of culture (mean ± SEM, n=3). (**g**) Fractional enrichment of [^13^C_4_]βOHB carbon in acetylated (M+2) metabolites from control (Ctrl) and Bdh1KO CD8^+^ T cells after 24 h of culture (mean ± SEM, n=3). **(h)** MID of [^13^C_4_]βOHB carbon in palmitate for activated control (Ctrl) and Bdh1KO CD8^+^ T cells after 24 h of culture (mean ± SEM, n=3).

We next used SCOT-knockdown T cells to identify the metabolic pathways downstream of ketolysis in CD8^+^ T cells. Using the competitive [U-^13^C_6_]glucose and [U-2,4-^13^C_2_]βOHB labeling strategy outlined in **Fig. 2d**, we observed loss of βOHB incorporation into TCA cycle intermediates, as well as acetyl-CoA (M+1) and downstream M+1 acetylated metabolites (i.e., Ac-spermidine, Ac-methionine) in *shOxct1*-expressing T cells (**Fig. S9**), mechanistically linking ketolysis to acetyl-CoA production in T cells. To further characterize the metabolic substrates contributing to acetyl-CoA production, we cultured activated CD8^+^ T cells in the presence of competitive uniformly ^13^C-labeled fuels at their physiologic concentrations (as in **Fig. 2b**) (*32*). Strikingly, over 50% of the intracellular acetyl-CoA (M+2) pool was derived from [U-^13^C_4_]βOHB, compared to less than 10% from [U-^13^C_6_]-glucose, in CD8^+^ T cells (**Fig. 3f**). The production of acetyl-CoA from βOHB was dependent on cell-intrinsic ketolysis, as [U-^13^C_4_]βOHB-dependent acetyl-CoA M+2 synthesis was decreased in both Bdh1KO and ketolysis-deficient (Bdh1KO plus SCOT-knockdown) T cells (**Figs. 3f**, **S10a).**

Further analysis revealed that [U-^13^C_4_]βOHB-dependent acetylation of intracellular metabolites, including Ac-carnitine, were significantly reduced in both Bdh1KO (**Fig. 3g**) and SCOT-knockdown (**Fig. S10b**) CD8^+^ T cells. We also observed significant levels of [U-^13^C_4_]βOHB-derived Ac-carnitine (M+2) in CD8^+^ T cells from *Lm*-OVA-infected animals infused with [U-^13^C_4_]βOHB at 6 dpi (**Fig. S10c**), providing evidence that βOHB is used to generate acetyl-CoA in T cells *in vivo*. Finally, we examined whether βOHB was used for *de novo* fatty acid synthesis, one of the critical biosynthetic pathways supported by cytosolic acetyl-CoA in proliferating cells (*45, 46*). Consistent with the high contribution of βOHB to the acetyl-CoA pool in T cells (**Fig. 3f**), we found that [U-^13^C_4_]βOHB carbon was incorporated into the fatty acid palmitate in a BDH1-dependent manner (**Fig. 3h**) and that [U-^13^C_4_]βOHB was preferred over [U-^13^C_6_]glucose for lipid synthesis in T cells (**Fig. S11**). Collectively, these results establish βOHB as a major, and previously unappreciated, substrate for acetyl-CoA synthesis in CD8^+^ T cells.

We next investigated the mechanistic impact of βOHB-dependent acetyl-CoA production on CD8^+^ Teff cell function. Consistent with the effect of βOHB boosting IFN-γ production by CD8^+^ T cells (**Figs. 1e-f**), we found that βOHB-treated CD8^+^ T cells displayed increased IFN-γ mRNA compared to controls, which required expression of BDH1 (**Fig. 4a**). Ketolysis-deficient CD8^+^ T cells cultured *in vitro* displayed reduced expression of mRNAs coding for IFN-γ and the cytotoxic protein granzyme B (GzmB) (**Fig. 4b**). Similarly, ketolysis-deficient CD8^+^ Teff cells responding to *Lm*-OVA at 7 dpi displayed reduced expression of several effector-associated genes (i.e., *Gzmk*, *Gzma*, *Cx3cr1, Klrg1*) (**Fig. 4c**), correlating with their reduced cytotoxicity signature *in vivo* (**Fig. 1i**). Together, these data indicate that KBs impact Teff cell function in part through transcriptional regulation of effector genes.

**Figure 4.**
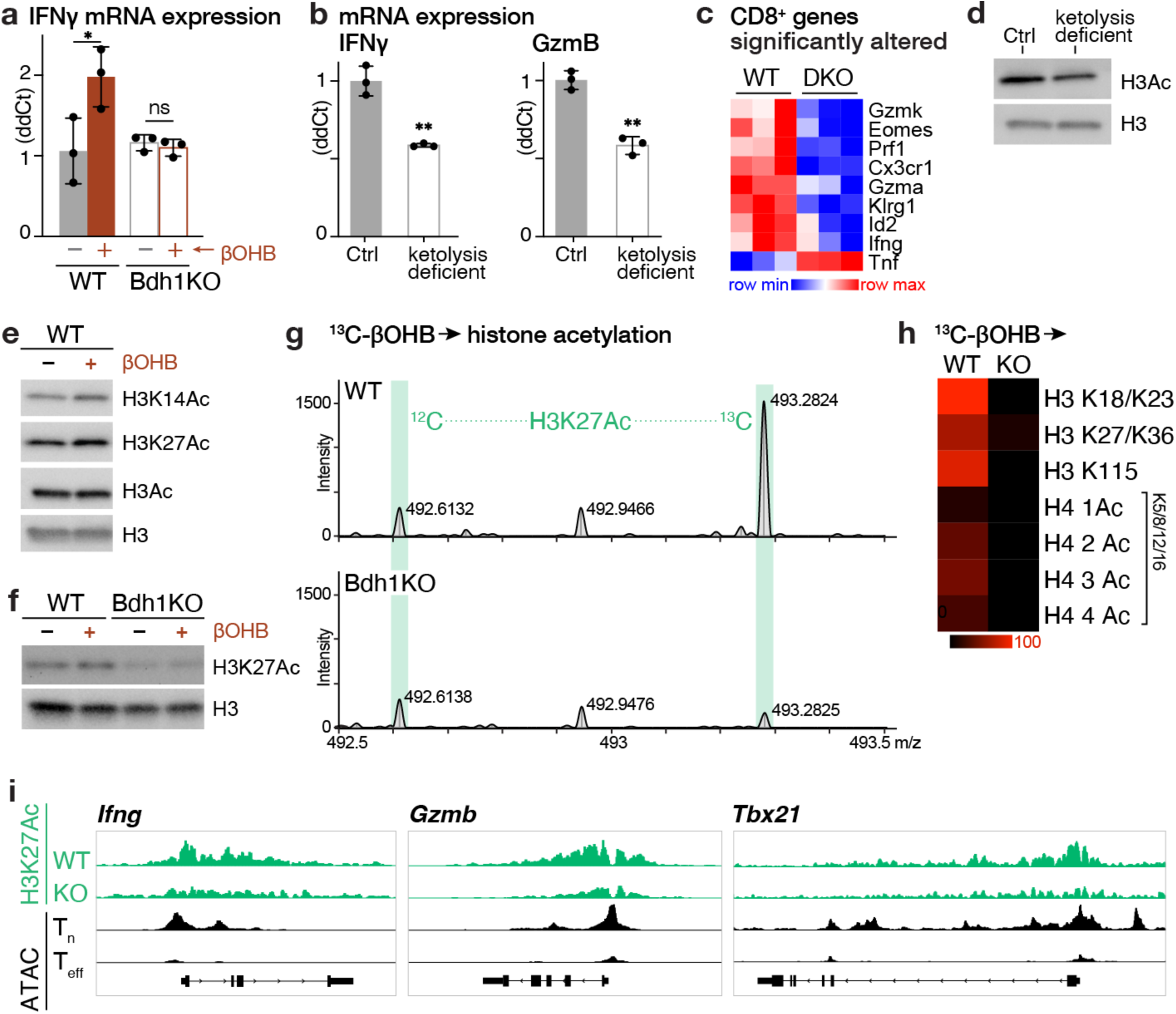
Ketolysis promotes histone acetylation of CD8^+^ T cell effector genes. (**a**) Relative abundance of *Ifng* mRNA transcripts in activated wild type (WT) and Bdh1KO CD8^+^ T cells cultured for 3 days in the presence (+) or absence (-) of 5 mM βOHB (mean ± SEM, n=3/group). (**b**) Relative abundance of *Ifng* and *Gzmb* mRNA transcripts in activated control (Ctrl) and ketolysis-deficient OT-I CD8^+^ T cells (mean ± SEM, n=3). (**c**) Heatmap of differentially expressed effector gene transcripts (*p<0.05*) altered in control (WT) and ketolysis-deficient (DKO) OT-I CD8^+^ cells isolated from *Lm*-OVA-infected mice 7 dpi (n=3/group). (**d**) Immunoblot of acetylated histone H3 in lysates from activated control (Ctrl) and ketolysis-deficient OT-I CD8^+^ cells. Total H3 levels are shown as a control for protein loading. **(e)** Immunoblot of global histone H3 acetylation (H3Ac) and specific acetylation at Lys14 (H3K14Ac) and Lys27 (H3K27Ac) in activated control (WT) T cells. **(f)** Immunoblot for H3 and H3K27Ac levels in activated control (WT) versus Bdh1KO CD8^+^ cells cultured for 24 h with (+) or without (-) 5 mM βOHB. (**g**) Representative MS spectra of histone H3 (peptide 27-40) from activated control (WT) or Bdh1KO CD8^+^ T cells cultured with [^13^C_4_]βOHB for 24 h. Peaks corresponding to unlabelled (^12^C) and ^13^C-labeled H3K27Ac peptides are highlighted in green. **(h)** Heatmap quantifying levels of [^13^C_4_]βOHB-dependent acetylation in histones H3 and H4 from activated control (WT) or Bdh1KO CD8^+^ T cells cultured with [^13^C_4_]βOHB for 24 h. For histone H4, the number of acetylated lysine residues on peptides containing Lys5/8/12/16 are quantified individually. **(i)** Data tracks for H3K27Ac peak enrichment at *Ifng*, *Gzmb*, and *Tbx21* gene loci for control (WT) and Bdh1KO CD8^+^ T cells activated for 24 h with plate-bound anti-CD3 and anti-CD28 antibodies. H3K27Ac peak enrichment is shown in green, with ATAC-seq tracks highlighting regions of chromatin accessibility in *in vivo* Tn and Teff cells (from (*22*)) shown in black. The data are representative of duplicate samples. *, *p* < 0.05; **, *p* < 0.01.

Chromatin remodeling following T cell activation helps to stabilize effector gene expression and reinforce CD8^+^ T cell effector functions (*18, 24*). Histone lysine (K) acetylation−specifically acetylation of lysine 27 on histone H3 (H3K27Ac)−is associated with chromatin accessibility and transcriptional activation (*47*), and is important for effector gene expression by CD8^+^ T cells (*48*). By immunoblotting of isolated histones, we observed reduced global histone H3 lysine acetylation in activated ketolysis-deficient CD8^+^ T cells (**Fig. 4d**). In addition, treatment of *in vitro*-activated CD8^+^ T cells with βOHB (5mM) was sufficient to globally increase histone H3 acetylation at several lysine residues compared to controls, including H3K14Ac and H3K27Ac (**Fig. 4e, S12**). βOHB has been implicated in epigenetic programming through its activity as a Class I histone deacetylase (HDAC) inhibitor or through direct β-hydroxybutylation of histones (*49, 50*). However, the effect of βOHB on H3K27Ac levels was reduced in Bdh1KO T cells (**Fig. 4f**), suggesting that ketolysis was directly driving changes in histone acetylation.

Given the major contribution of βOHB to the acetyl-CoA pool in T cells (**Fig. 3f**), we hypothesized that βOHB-derived acetyl-CoA was being used to acetylate histones. To test this directly, we cultured *in vitro* activated CD8^+^ T cells with [U-^13^C_4_]βOHB and measured the enrichment of ^13^C-labelled acetyl-groups on histone H3 via mass spectrometry (**Fig. 4g-h**). Carbon from [U-^13^C_4_]βOHB was enriched on histone H3K27 in control but not BDH1-deficient T cells (**Fig. 4g**), establishing ketolysis as an obligate step for βOHB-dependent H3K27 acetylation in CD8^+^ T cells. We also observed enrichment of [U-^13^C_4_]βOHB carbon in acetyl groups from multiple lysine residues on histone H3 (K18, K23, K36, K115) and H4 (K5, K8, K12, K16) in control but not BDH1-deficient T cells (**Fig. 4h**). Next, we used quantitative chromatin immunoprecipitation coupled to next generation sequencing (ChIP-seq) (*51*) to assess the impact of BDH1-deficiency on H3K27Ac modifications in T cells at genome-scale resolution. Overall, we observed a global reduction of H3K27Ac levels in activated BDH1-knockout T cells compared to controls (**Fig. S13a**). Notably, we observed reduced levels of H3K27Ac at the promoters of effector gene loci−including *Ifng*, *Gzmb*, *Prf1*, and *Tbx21* (Tbet)−in BDH1-deficient T cells compared to controls (**Fig. 4i, S13b**). Interestingly, H3K27Ac enrichment at genetic loci associated with CD8^+^ T cell identity and signal transduction (i.e., *Cd3e, Cd8a, Zap70*) were similar between control and BDH1-deficient T cells (**Fig. S13b**). Collectively, we conclude from these data that KB-dependent acetyl-CoA production supports CD8^+^ T cell effector function in part by directly altering acetylation-dependent epigenetic programming at effector gene loci.

Infection-induced anorexia and increased KB production are host metabolic adaptations to pathogen challenge with poorly defined impact on immune responses. Here, we identify cell-intrinsic ketolysis as a non-redundant metabolic feature of CD8^+^ T cells that boosts effector function by altering T cell metabolic and epigenetic programming. Our data establish KBs as potent bioenergetic substrates for CD8^+^ T cells−preferred over glucose for oxidation−that increase mitochondrial respiratory capacity and fuel TCA cycle-dependent biosynthesis both *in vitro* and *in vivo*. We speculate that ketolysis evolved as a metabolic program to preserve T cell effector function during periods of starvation or inflammation-driven changes in feeding behavior (*52*). While immunomodulatory effects of KBs on immune cell populations have been documented (*17, 35, 53–55*), our results point towards ketolysis-driven acetyl-CoA synthesis as a key mechanism underlying the immunomodulatory effects βOHB on T cell function. We establish that βOHB is a major source of acetyl-CoA production in CD8^+^ T cells, which directly contributes to histone acetylation, and promotes permissive H3K27 acetylation at effector gene loci (i.e., *Ifng, Gzmb, Tbx21*) in activated CD8^+^ T cells. Our data highlight ketolysis as a potential target for advancing T cell-based immunotherapies. Such therapeutic manipulation could be achieved by altering systemic concentrations of circulating KBs through the diet (i.e., ketogenic diet, ketone ester supplementation) (*56, 57*), and may benefit CD8^+^ T cell function when conventional fuels such as glucose are limiting.

## Acknowledgements

**General:** We acknowledge Drs. Ralph DeBerardinis, Julian Lum, Sara Nowinski, and members of the Jones and Krawczyk laboratories for scientific discussions contributing to this manuscript. We thank Teresa Leone, Matthew Vos, Jeanie Wedberg, and Michelle Minard for administrative assistance. We thank members of the VAI Core Facilities (Metabolomics and Bioenergetics, Genomics, Bioinformatics and Biostatistics, and Flow Cytometry) for technical assistance.

**Funding:** MJW is supported by a National Cancer Institute (NCI) T32 training grant (T32CA251066-01A1). JL is supported by a Van Andel Institute (VAI) Metabolism & Nutrition (MeNu) Program Pathway-to-Independence Award and Canadian Institutes of Health Research (CIHR) Fellowship (MFE-181903). DPK is supported by the National Institutes of Health (NIH, R01HL128349 and R01HL151345). CMK is supported by the National Institute of Allergy and Infectious Diseases (NIAID, R21AI153997) and VAI. FMB is a FRQS Senior scholar (award number 281824) and is supported by the National Sciences and Engineering Research Council of Canada (RGPIN-2018-05414). SBR is supported by the National Institute of General Medical Sciences (NIGMS, R35GM124736) and VAI. PAC and PP are supported by the National Institute of Diabetes and Digestive and Kidney Diseases (NIDDK, DK091538) and National Institute on Aging (NIA, AG069781). RGJ is supported by the Paul G. Allen Frontiers Group Distinguished Investigator Program, NIAID (R01AI165722), and VAI.

**Author Contributions:** Conceptualization, KML and RGJ; Experimental Design, KML, EHM, DPK, PP, CMK, FMB, RDS, SBR, PAC, and RGJ; Investigation, KML, SMKG, EHM, MJW, JL, LRD, BMO, AK, IK, LMD; Data Analysis, KML, SKMG, EHM, MJW, JL, ZF, ZM, BMO, BMD, MJW, BF, KSW, RDS, and RGJ; Writing – Original Draft, KML, KSW, and RGJ; Writing – Editing, KML, CMK, SBR, PAC, and RGJ; Visualization, ZF and KSW; Supervision, RGJ; Funding Acquisition, RGJ.

**Competing Interests**: RGJ is a scientific advisor for Agios Pharmaceuticals and Servier Pharmaceuticals and is a member of the Scientific Advisory Board of Immunomet Therapeutics.

## Data and Materials Availability

RNA sequencing data used for meta analysis in Figure 1 are available at NCBI GEO (accession numbers (GSE86881, GSE89307, GSE84820)). RNA sequencing data on ketolysis-deficient T cells from **Figure 1** are available at NCBI GEO (accession: GSE212048; token: clepicoczzcffop). The mass spectrometry proteomics data have been deposited to the ProteomeXchange Consortium via the PRIDE partner repository with the dataset identifier PXD036292 (access via username; password: sKA2ueMn). Bioenergetics data analysis was based on protocols developed by Mookerjee and Brand (*58*) which is available for download at https://russelljoneslab.vai.org/tools. All the data are available in the main text or supplemental figures and tables. Additional information and request for resources and reagents should be directed to and will be made available by the corresponding author, Russell G. Jones.

## Supplementary Materials

## Materials and Methods

### Mice

C57BL/6, CD90.1 (Thy1.1^+^), B6.SJL-*Ptprc^a^ Pepc^b^*/BoyJ, Tg(TcraTcrb)1100Mjb (OT-I), and *Cd4-Cre* mice were purchased from The Jackson Laboratory. *Bdh1*-floxed animals were generated by Daniel Kelly (*1*). Mice were bred and maintained under specific pathogen-free conditions at VAI under approved protocols. Genotyping was performed using tail biopsies using defined primer sets (see Table S4). Experiments were performed using mice (of both sexes) between 8 and 20 weeks of age.

### T cell purification and culture

For mouse T cell isolation, naïve CD8^+^ T cells were purified from spleen and peripheral lymph nodes by negative selection (StemCell Technologies, Vancouver, BC) as previously described (*2*). Cells were cultured in Iscove’s Modified Delbecco’s Medium (IMDM) or Van Andel Institute-modified Iscove’s Medium (VIM) (*3*) supplemented with 10% dialyzed FBS (Wisent, St. Bruno, QC), penicillin-streptomycin (Invitrogen), and 2-ME (Sigma-Aldrich, St. Louis, MO). Unless stated otherwise, CD8^+^ T effector (Teff) cells were generated by activation in standard IMDM containing 25 mM glucose and 6 mM L-glutamine. For physiologic culture conditions, Seahorse experiments, and ^13^C tracing experiments, CD8^+^ Teff cells were culture in medium containing 5 mM glucose and 0.5 mM L-glutamine, respectively. Physiologic carbon sources (PCS) were added to culture medium (IMDM or VIM) as indicated at the following concentrations: acetate (400 μM), βOHB (850 μM), citrate (215 μM), lactate (3 mM), and pyruvate (150 μM) (*3*). *In vitro*-activated CD8^+^ Teff cells were generated by stimulating naïve CD8^+^ T cells (1 x 10^6^ cells/mL) with plate-bound anti-CD3ε (clone 2C11) and anti-CD28 (clone 37.51) antibodies (eBioscience, San Diego, CA) for 3 days. For extracellular flux analysis, stable isotope labelling, and histone isolation procedures activated CD8^+^ Teff cells were re-cultured (5 × 10^6^ cells/well in 6-well plates) for up to 48h in IMDM containing 50 U/mL IL-2 (PeproTech, Rocky Hill, NJ) prior to use in assays. For retroviral transduction experiments, CD8^+^ Thy1.1^+^ OT-I T cells (wild type or *Bdh1-*deficient) were transduced with retrovirus 24 hours post activation and expanded for 2 additional days in IMDM containing IL-2 as previously described (*2, 4*). Transduced T cells were FACS sorted and cultured overnight prior to adoptive transfer into naïve Thy1.2^+^ C57BL/6 hosts.

### Adoptive transfer and infection with *L. monocytogenes* (*Lm*-OVA)

Mice were immunized intravenously with a sublethal dose of recombinant attenuated Listeria monocytogenes expressing OVA (*Lm*-OVA, 2 x 10^6^ CFU) as previously described (*5, 6*). For adoptive transfer experiments 6-7 days post infection (dpi) using non-transduced or transduced cells, 5 x 10^3^ CD8^+^ OT-I T cells (Thy1.1^+^ or CD45.2^+^) were injected intravenously into C57BL/6 mice (Thy1.2^+^CD45.2^+^ or CD45.1^+^), followed by *Lm*-OVA infection 1 day later. Splenocytes were isolated from mice 7 dpi and analyzed for the presence of OVA-specific CD8^+^ T cells by Thy1.1 or CD45.2 staining and cytokine production analyzed by intracellular cytokine staining (ICS) following peptide re-stimulation (OVA_257-264_) as previously described (*5, 6*). For metabolic analysis of *Lm*-OVA-specific Thy1.1^+^ OT-I T cells *in vivo* using ^13^C-labeled metabolites, Thy1.2^+^ C57BL/6 mice received 2.5 x 10^6^ or 5 x 10^4^ Thy1.1^+^ OT-I T cells for analysis at 2 and 6 dpi, respectively. *Lm*-OVA-specific CD8^+^ OT-I T cells were isolated from the spleen of infected mice by positive selection using the EasySep mouse CD90.1 positive selection kit (StemCell Technologies) as previously described (*2, 7*).

### Flow cytometry

Single cell suspensions from the spleen were surface stained with a cocktail of fluorescently labelled antibodies listed in Table S5. Cell viability was assessed by using Fixable Viability Dye eFluor 780 (eBioscience) according to manufacturer’s protocols. To assess cytokine production, splenocytes were plated in the presence of PMA (50ng/ml) and Ionomycin (50ng/ml and Merck) for 2h and with Brefeldin A (5ug/ml, Biolegend) added for the last 2h of stimulation prior surface staining. After restimulation, cells were surface stained, fixed, and permeabilized using FoxP3/Transcription Factor Staining Buffer Set (eBioscience), followed by processing for intracellular staining using fluorescently labelled antibodies. For analysis of antigen-specific response to *Lm*OVA, splenocytes harvested 7 dpi were stimulated with OVA257-264 peptide using previously published protocols (*5, 6*). Flow cytometry was performed on Cytoflex (Beckman Coulter) or Aurora Cytek cytometers and cell sorting on Astrios (Beckman Coulter) or BD FACSAria Fusion cell sorters. Data analysis was performed using FlowJo software (Tree Star).

### Extracellular flux analysis

T cell oxygen consumption rate (OCR) and extracellular acidification rate (ECAR) were measured using a Seahorse XF96 Extracellular Flux Analyzer following established protocols (*2*). Activated and IL-2-expanded T cells (1.5 x 10^5^/well) were cultured in XF medium containing 5mM glucose and 0.5mM glutamine, following centrifugation onto poly-D-lysine-coated XF96 plates, and cellular bioenergetics assessed at 5 minute intervals following the sequential addition of oligomycin (2.0 μM), fluoro-carbonyl cyanide phenylhydrazone (FCCP, 2.0 μM), rotenone/antimycin A (2 μM), and monensin (10mM). Data were normalized to cell number. Where indicated, βOHB (2 mM) and AcAc (2 mM) were added to cell culture for 2 h prior to Seahorse analysis and added to Seahorse medium over the assay period. Bioenergetics data analysis was based on protocols developed by Mookerjee and Brand (*8*), which is available for download at https://russelljoneslab.vai.org/tools.

### Acetoacetate synthesis

[^12^C_4_]-ethyl acetoacetate (Merck) and [^13^C_4_]-ethyl-acetoacetate (Cambridge Isotope labs) were hydrolysed by mixing 1ml of ethyl acetoacetate with 8 ml 1M NaOH at 60°C by stirring for 30 min. Hydrolyzed samples were placed on ice and adjusted to pH 7.5 using 50% HCl. Subsequently, samples were frozen at -80°C. The concentration of synthesized acetoacetate was evaluated using the Autokit Total Ketone Bodies Assay (Fujifilm Wako) and used for *in vitro* assays.

### Stable isotope labeling (SIL) and metabolomics

SIL experiments with *in vitro*-activated T cells using liquid chromatography (LC) or gas chromatography (GC) coupled to mass spectrometry (MS) were conducted as previously described (*2, 3*). In brief, naïve CD8^+^ T cells were activated as above washed, in IMDM or VIM containing 10% dialyzed FBS, and re-cultured (2.5 × 10^6^ cells/well in 24-well plates) for indicated times in medium containing ^13^C-labeled metabolites (Cambridge Isotope Laboratories) at the following concentrations: [U-^13^C_6_]glucose, 5 mM; [U-^13^C_2_]acetate, 400 μM; [U-^13^C_4_]AcAc, 2 mM; [U-^13^C_3_]alanine, 200 μM; [U-^13^C_4_]βOHB, 0.85 or 2 mM; [^13^C_6_]citrate, 215 μM; [^13^C_3_]lactate, 3 mM; [^13^C_3_]pyruvate, 150 μM. Cells were transferred from tissue culture plates to falcon tubes and centrifuged at 500g 4°C for 3 min. The cell pellet was washed with ice-cold saline and centrifuged before being snap frozen on dry-ice and stored at -80°C. Metabolites were extracted by modified Bligh-Dyer extraction (*9*) by the addition of ice-cold methanol (A456, Fisher Scientific) directly to frozen cells, to which one volume of chloroform (A456, Fisher Scientific) was added. The sample was vortexed for 10 s, incubated on ice for 30 min, and then 0.9 parts of LC-MS grade water (W6-4, Thermo Fisher Scientific) was added. The samples were vortexed vigorously and centrifuged at maximum speed to achieve phase separation. The top layer containing polar metabolites was aliquoted into a fresh tube and dried in a speedvac for LC-MS analysis. The bottom layer was retained for fatty acid methyl-ester measurement.

For LC-MS analysis, metabolite extracts were resuspended in 50 µL of 60% acetonitrile (A955, Fisher Scientific) and analyzed by high resolution accurate mass spectrometry using an ID-X Orbitrap mass spectrometer (Thermo Fisher Scientific) coupled to a Thermo Vanquish Horizon liquid chromatography system. 2 µL of sample volume was injected on column. Chromatographic separations were accomplished with Acquity BEH Amide (1.7 µm, 2.1 mm x 150 mm) analytical columns (#176001909, Waters, Eschborn, Germany) fitted with a pre-guard column (1.7 µm, 2.1mm x 5 mm; #186004799, Waters) using an elution gradient with a binary solvent system. Solvent A consisted of LC/MS grade water (W6-4, Fisher), and Solvent B was 90% LC/MS grade acetonitrile (A955, Fisher). For negative mode analysis, both mobile phases contained 10 mM ammonium acetate (A11450, Fisher Scientific), 0.1% (v/v) ammonium hydroxide, and 5 µM medronic acid (5191-4506, Agilent Technologies). For positive mode analysis, both mobile phases contained 10 mM ammonium formate (A11550, Fisher), and 0.1% (v/v) formic acid (A11710X1, Fisher). For both negative and positive mode analyses the 20-min analytical gradient at a flow rate of 400 µL/min was: 0–1.0 min ramp from 100% B to 90% B, 1.0–12.5 min from 90% B to 75% B, 12.5–19 min from 75% B to 60% B, and 19–20 min hold at 60% B. Following every analytical separation, the column was re-equilibrated for 20 min as follows: 0–1 min hold at 65%B at 400 µL/min, 1–3 min hold at 65% B and ramp from 400 µL/min to 800 µL/min, 3–14 min hold at 65% B and 800 µL/min, 14–14.5 min ramp from 65% B to 100% B at 800 µL/min, 14.5–16 min hold at 100% B and increase flow from 800 µL/min to 1200 µL/min, 16–18.4 min hold at 100% B at 1200 µL/min, 18.4–19.5 min hold at 100% B and decrease flow from 1200 µL to 400 µL/min, 19.5–20 min hold at 100% B and 400 µL/min. The column temperature was maintained at 40°C. The H-ESI source was operated at spray voltage of 2500 V(negative mode)/3500 V(positive mode), sheath gas: 60 a.u., aux gas: 19 a.u., sweep gas: 1 a.u., ion transfer tube: 300°C, vaporizer: 300°C. For isotopically labelled experimental replicates, high resolution MS^1^ data was collected with a 20-min full-scan method with m/z scan range using quadrupole isolation from 70 to 1000, mass resolution of 120,000 FWHM, RF lens at 35%, and standard automatic gain control (AGC). Unlabelled control samples were used for data dependent MS^2^ (ddMS^2^) fragmentation for compound identification and annotation via the AquireX workflow (Thermo Scientific). In this workflow, first blank and experimental samples are injected to generate exclusion and inclusion lists, respectively, followed by iterative sample injections for ddMS^2^ fragmentation where triggered ions are added to the exclusion list for subsequent injections. ddMS^2^ data was collected using MS1 resolution at 60,000, MS^2^ resolution at 30,000, intensity threshold at 2.0 x 10^4^, and dynamic exclusion after one trigger for 10 s. MS^2^ fragmentation was completed first with HCD using stepped collision energies at 20, 35, and 50% and was followed on the next scan by CID fragmentation in assisted collision energy mode at 15, 30, and 45% with an activation Q of 0.25. Both MS^2^ scans used standard AGC and a maximum injection time of 54ms. The total cycle time of the MS^1^ and ddMS^2^ scans was 0.6 s.

Full scan LC-MS data were analyzed in Compound Discoverer (v 3.2, Thermo Scientific). Compounds were identified by chromatography specific retention time of external standards and MS^2^ spectral matching using the mzCloud database (Thermo Scientific).

TCA cycle intermediates were measured via GC-MS following LC-MS analysis. Briefly, following LC/MS extracts were dried and derivatized with 30 µL of methoxyamine (11.4 mg/mL) in pyridine and 70 µL of MTBSFA+1%TMCS as described previously (*10*). In addition, GC-MS was used to evaluate incorporation of 13C into de novo synthesized fatty acids from metabolic precursors using fatty acid methyl-esters (FAMEs). The bottom organic fraction from the Bligh-Dyer extraction (above) was aliquoted, dried in a speedvac, and FAMEs were generated as described previously (*11*). GC-MS analysis of both TBDMS derivatives and FAMEs were conducted on an Agilent 7890/5977b GC/MSD equipped with a DB-5MS+DG (30 m x 250 µm x 0.25 µm) capillary column (Agilent J&W, Santa Clara, CA, USA) was used. Data were collected by electron impact set at 70 eV. A total of 1 μL of the derivatized sample was injected in the GC in split mode (1:2 or 1:4) with inlet temperature set to 280°C, using helium as a carrier gas with a column flow rate of 1.2 mL/min. The oven program for all metabolite analyses started at 95°C for 1 min, increased at a rate of 40°C/min until 118°C and held for 2 min, then increased to 250°C at a rate of 12°C/min, then increased to 320°C at a rate of 40°C/min and finally held at 320°C for 7 min. The source temperature was 230°C, the quadrupole temperature was 150°C, and the GC-MS interface at 285°C. Data were acquired both in scan mode (50–800 m/z) and 2 Hz.

MassHunter software (v10, Agilent Technologies) was used for peak picking and integration of GC-MS data. Peak areas of all isotopologues for a molecular ion of each compound in both labeled experimental and unlabeled control samples were used for mass isotopologue distribution analysis via a custom algorithm developed at VAI. This algorithm uses matrices correcting for natural contribution of isotopologue enrichment were generated for each metabolite as described previously (*12, 13*).

### *In vivo* ^13^C tracer infusions

*In vivo* infusions of *Lm*OVA-infected mice were conducted using previously described protocols (*2, 7*). Briefly, mice were anesthetized using isoflurane and infused intravenously with ^13^C tracers over a 2 h period. [U-^13^C_6_]glucose infusions were performed as previously described, using a stock concentration of 100 mg/ml, an initial bolus dose of 120 μL and infusion rate of 2.5 μL/min for 2 h for a 20 g mouse. For infusions of sodium D-[U-^13^C_4_]βOHB, mice received an initial bolus of 0.487 mg per gram of mouse, followed by infusion of 0.05 μL/min/g bodyweight using a 1.5 M stock concentration for 2 hours. Subsequently, mice were cervically dislocated, plasma was harvested via cardiac puncture, centrifuged at 500g 4°C for 3 min, and snap frozen in liquid nitrogen. Spleens were collected and processed for Thy1.1^+^ T cell isolation by magnetic bead isolation as previously described (*7*). Isolated T cells were snap frozen on dry ice and subsequently processed for metabolomic analysis as described.

### RNA isolation, sequencing, and qPCR analysis

Total RNA was isolated from murine T cells via RNeasy Kit (Qiagen) with DNase digestion (Qiagen) following manufacturer’s instructions. For quantitative PCR (qPCR) analysis, total RNA was reverse transcribed using a High Capacity cDNA Reverse Transcriptase kit (Life Technologies) and qPCR performed using SYBR green (Bio-Rad). Results were normalized to *Stk11* mRNA levels and wild type controls using standard ddCt methods. RNA preparation and library construction for RNA sequencing was conducted by the VAI Genomics Core as previously described (*14*). Libraries were sequenced using a NovaSeq 6000 (Illumina) using 50 bp paired-end sequencing (5x10^7^ reads/sample). Gene-set enrichment analysis (GSEA) on RNA-seq data was conducted using the gage function and non-parametric Kolmogorov–Smirnov test from the GAGE (version 2.22.0) R Bioconductor package (*15*). RNA-seq data files are available at NCBI GEO (accession: GSE212048).

For the meta-analysis in **Figure 1**, raw sequences from RNA-sequencing of CD8^+^ T cells from three previously published studies (GEO accessions: GSE89307, GSE84820, and GSE86881) were downloaded. Adaptor sequences and low-quality reads were trimmed using Trim Galore (v0.6.0) (*16*). Trimmed reads were aligned to the mm10 reference genome using STAR (v2.7.8) (*17*). Count tables of all samples were then imported into limma (v3.48.3) (*18*), with a batch variable included in the design matrix to account for the different study designs and sequencing platforms. Pearson correlations among samples were calculated using batch corrected, variance stabilization transformed counts using DESeq2 (v1.32) (*19*), and the Pearson correlation matrix was used to generate a heatmap. Batch corrected counts were also used to conduct principal component analysis (PCA) using DESeq2. Differential gene expression analyses of two main comparisons (comparison 1: cancer Tex cells vs. Teff cells; comparison 2: virus Tex cells vs. Teff cells) were conducted on raw counts using DESeq2, with a covariate to adjust for batch, and Benjamini-Hochberg adjusted p-values to maintain a 5% false discovery rate. The median of the two Wald test statistics for each gene, one from each of the two comparisons, were used for gene rankings. Gene ontology (GO) terms for each gene was retrieved from BioMart using R package biomaRt (v2.48.4) (*20, 21*). Lastly, for each comparison, genes were sorted by their fold change from highest to lowest (using Teff cells as reference in both comparisons) and gene set enrichment analysis (GSEA) (*22*) was conducted for each comparison independently using clusterProfiler (v4.0.5) (*23*). Gene sets C2, C5, C6, C7, and Hallmark for mm10 was retrieved from Molecular Signatures Database using R package MSigDB (7.4.1).

### Immunoblotting

Cells were lysed in modified Laemmli lysis buffer (240 mM Tris/HCl pH 6.8, 40% glycerol, 8% SDS, 5% b-ME) supplemented with protease and phosphatase inhibitors (Roche/Sigma-Aldrich). A Pierce BCA Protein Assay Kit (Thermo Fisher Scientific, Waltham, MA, USA) was used to quantify protein from whole cell lysates. Lysates were resolved by SDS-PAGE, transferred to nitrocellulose, and incubated with primary antibodies to Bdh1, Oxct1, a-tubulin or β-actin, and HRP-conjugated secondary antibodies. Histone proteins were extracted from T cells using a Histone Extraction Kit (Abcam) following manufacturer’s instructions. Lysates containing 0.5-2 μg of protein were resolved on a 4-20% SDS-PAGE gel and transferred to PVDF membrane. Membranes were incubated with primary antibodies overnight at 4°C, and then incubated with secondary antibody for 1hr at room temperature prior to exposure. Primary antibodies are listed in Table S5.

### Proteomic analysis of histone proteins

For proteomic analysis of histones, activated CD8^+^ T cells were cultured in VIM medium (5 mM glucose, 0.5 mM glutamine) containing 5 mM standard [^12^C_4_]βOHB or [^13^C_4_]βOHB for 24 h. Histone isolation was carried out using the same methodology for immunoblotting, with the addition of 2 ice-cold PBS washes. Histone extracts were lyophilized and resuspended in 50 μL of 8M urea with 10 mM HEPES-KOH pH 7.5. Proteins were reduced by adding dithiothreitol (DTT) to a final concentration of 5 mM and by heating at 95°C for 2 minutes, followed by a 30 minute incubation at room temperature. The alkylation of the proteins was carried out by adding chloroacetamide (Sigma-Aldrich, Saint-Louis) to a final concentration of 7.5 mM followed by a 20 minute incubation in the dark at room temperature. The urea concentration was then diluted to a final concentration of 2 M by adding 150 μL of 50 mM ammonium bicarbonate (NH_4_HCO_3_) (Sigma-Aldrich, Saint-Louis). Digestion was started by adding 1 μg of Pierce MS-grade trypsin (Thermo Fisher Scientific, Waltham) and incubated overnight at 30°C with shaking. Sample was then acidified by adding Trifluoroacetic acid (TFA) (Sigma-Aldrich, Saint-Louis) to a final concentration of 0.2%.

The peptides were purified with ZipTip 100-μl micropipette tips containing a C18 column, according to the manufacturer’s protocol (EMD Millipore, Burlington, VT) and eluted with 300 μL of 50% ACN/1% FA buffer in a new low-binding microtube. The peptides were then concentrated by centrifugal evaporator at 60°C until complete drying (∼3 h) and then resuspended in 50 μL of 1% FA buffer. Peptides were assayed using a NanoDrop spectrophotometer (Thermo Fisher Scientific, Waltham, MA) and read at an absorbance of 205 nm. The peptides were then transferred to a glass vial (Thermo Fisher Scientific) and stored at −20 °C until analysis by mass spectrometry.

For LC-MS analysis, 250 ng of peptides were injected into an HPLC (nanoElute, Bruker Daltonics) and loaded onto a trap column (Acclaim PepMap100 C18 column, 0.3 mm id x 5 mm, Dionex Corporation) with a constant flow of 4 µL/min, then eluted onto an analytical C18 Column (1.9 µm beads size, 75 µm x 25 cm, PepSep). Peptides were eluted over a 2-hour gradient of ACN (5-37%) in 0.1% FA at 400 nL/min while being injected into a TimsTOF Pro ion mobility mass spectrometer equipped with a Captive Spray nano electrospray source (Bruker Daltonics). Data was acquired using data-dependent auto-MS/MS with a 100-1700 m/z mass range, with PASEF enabled with several PASEF scans set at 10 (1.17 seconds duty cycle) and a dynamic exclusion of 0.4 min, m/z dependent isolation window and collision energy of 42.0 eV. The target intensity was set to 20,000, with an intensity threshold of 2,500.

Raw data files were analyzed using MaxQuant (version 2.0.3.0) and the Uniprot human proteome database (21/03/2020, 75,776 entries). The settings used for the MaxQuant analysis (with TIMS-DDA type in group-specific parameters) were: 4 miscleavages were allowed; the minimum peptide length was set to 5; enzyme was Trypsin (K/R not before P); fixed modification was carbamidomethylation on cysteine; variable modifications were methionine oxidation, protein N-terminal acetylation (^12^C, ^13^C), lysine (K) acetylation (^12^C, ^13^C) and protein carbamylation (K, N-terminal). A mass tolerance of 20 ppm was used for both precursor and fragment ions. Identification values "PSM FDR", "Protein FDR" and "Site decoy fraction" were set to 0.05. Minimum peptide count was set to 1. Both the "Second peptides" and "Match between runs" options were also allowed. Compass DataAnalysis (version 5.3, Bruker Daltonics) was further used to represent both spectral and graphical MS1 results for specific peptides.

### Chromatin immunoprecipitation sequencing (ChIP-seq) analysis

ChIP-seq analysis of H3K27Ac enrichment in CD8^+^ T cells was conducted using the sans spike-in quantitative ChIP (siQ-ChIP) method as previously described (*24, 25*). Naïve CD8^+^ T cells (5 x 10^6^) were activated with anti-CD3 and anti-CD28 antibodies for 24 h, then rinsed once with 10 mL of D-PBS (Gibco, 14190136) followed by cross-linking in suspension for 5 min in 10 mL of 0.75% formaldehyde (Pierce, 28906) in D-PBS at room temperature. Prior to every liquid removal step, cells were centrifuged at 800 RCF for 3 min. Formaldehyde was removed, and cells were quenched for 5 min with 10 mL of 750 mM Tris pH 10-10.5. Cells were washed twice with 10 mL of D-PBS, snap-frozen in liquid nitrogen, and stored -80°C. Cells were then lysed under hypotonic conditions (1 mL of 20 mM Tris-HCl pH 8, 85 mM KCl, 0.5% NP-40 (1 tablet of protease inhibitor (Roche, 11836170001) per 5 mL of buffer)) for 30 min on ice. Nuclei were collected by centrifugation at 1300 RCF for 5 min, lysed by resuspension in nuclei lysis buffer (150 μL 50 mM Tris-HCl pH 8,150 mM NaCl, 2 mM EDTA, 1% NP-40, 0.5% sodium deoxycholate, 0.1% SDS (1 tablet of protease inhibitor per 5 mL of buffer)), and passed through a 27-gauge needle (BD #309623 Lot: 0227218). Lysates were diluted to 500 μL by addition of binding buffer (25 mM HEPES pH 7.5, 100 mM NaCl, 0.1% NP-40). 5 μL of RNAse A/T1 (Thermo Scientific, EN0551) was added, and the sample was incubated at 37°C for 25 min. Next, CaCl_2_ was added to a final concentration of 40 mM, followed by the addition of 75 U of micrococcal nuclease (Worthington Biochemical) and incubated at 37°C for 5 min. MNase was quenched by the addition of 40 mM EDTA, and the total volume was brought to 1.2 mL with binding buffer. Next, insolubilities were removed by centrifugation at max speed (about 21,000 RCF) at 4°C for 5 min, and the supernatant containing soluble chromatin was collected. At this stage, 5 μL of chromatin was measured using the Qubit dsDNA HS Assay Kit (Invitrogen, Q32851).

Samples were diluted with binding buffer to ensure similar chromatin concentrations and to match IP conditions. 50 μL of chromatin was set aside for input for each sample. For each IP, 25 μL of Protein A coated magnetic beads (Invitrogen, 10008D) were washed once with binding buffer and incubated with either 0, 2.5, or 10 μL of H3K27Ac antibody (Active Motif #39133, lot: 16119013). Total volume of bead plus antibody was brought to 200 μL using binding buffer and were rotated at room temperature for 30 min. For comparison of PTM between samples, we performed one biological replicate of all samples on the same day and made a master mix of bead plus antibody, scaling up all components by the number of samples. For example, for 2 samples with technical replicate points, bead and antibody amounts were scaled by x 4.2 to account for dead volume. Buffer containing antibody was removed, and bead plus antibody were resuspended in 200 μL of soluble chromatin followed by 30 min rotation at room temperature. Unbound chromatin was removed, and beads were vortexed for 10 s with 500 μL of binding buffer. Buffer was removed, and bound material was eluted from beads by vortexing for 10 s in 133 μL of elution buffer (25 mM HEPES pH 7.5, 100 mM NaCl, 1% SDS, and 0.1% NP-40). At this time, the input was brought to 133 μL by the addition of 83 μL elution buffer. Proteinase K (Invitrogen, 25530015) was added to a final concentration of 15 μM and the sample was incubated overnight at 37°C. The following morning, each DNA sample was purified using MinElute PCR Kit (Qiagen, 28004) and eluted in 30 μL of Buffer EB. Five μL of DNA was quantified by Qubit dsDNA HS Assay Kit. The remaining 25 μL of DNA was frozen at - 20°C until library preparation.

Libraries were sequenced using a NextSeq 500 (Illumina) using 75 bp paired-end sequencing (5x10^7^ reads/sample for IPs, 1x10^8^ reads/sample for input). Next generation sequencing (NGS) data were aligned to the mm10 genome. Bed files generated by the NGS alignment were processed using the latest siQ-ChIP release (found at https://github.com/BradleyDickson/siQ-ChIP<u>) and</u> siQ-ChIP quantification performed as previously described (*24*). Responses were computed automatically by the siQ-ChIP software as the ratio of area under overlapping peaks for any pair of tracks being compared. Individual ChIP tracks were visualized using IGV (*26*). Input files for siQ-ChIP and resulting output files and gnuplot script for plotting will be published in NCBI GEO.

### Statistics

Statistical analysis was assessed by GraphPad Prism software (GraphPad) by Student’s t-test. Statistical significance is indicated in all figures by the following annotations: *, *p* < 0.05; **, *p* < 0.001; ***, *p* < 0.0001; ****, *p* < 0.00001.

**Figure S1, related to Figure 1.**
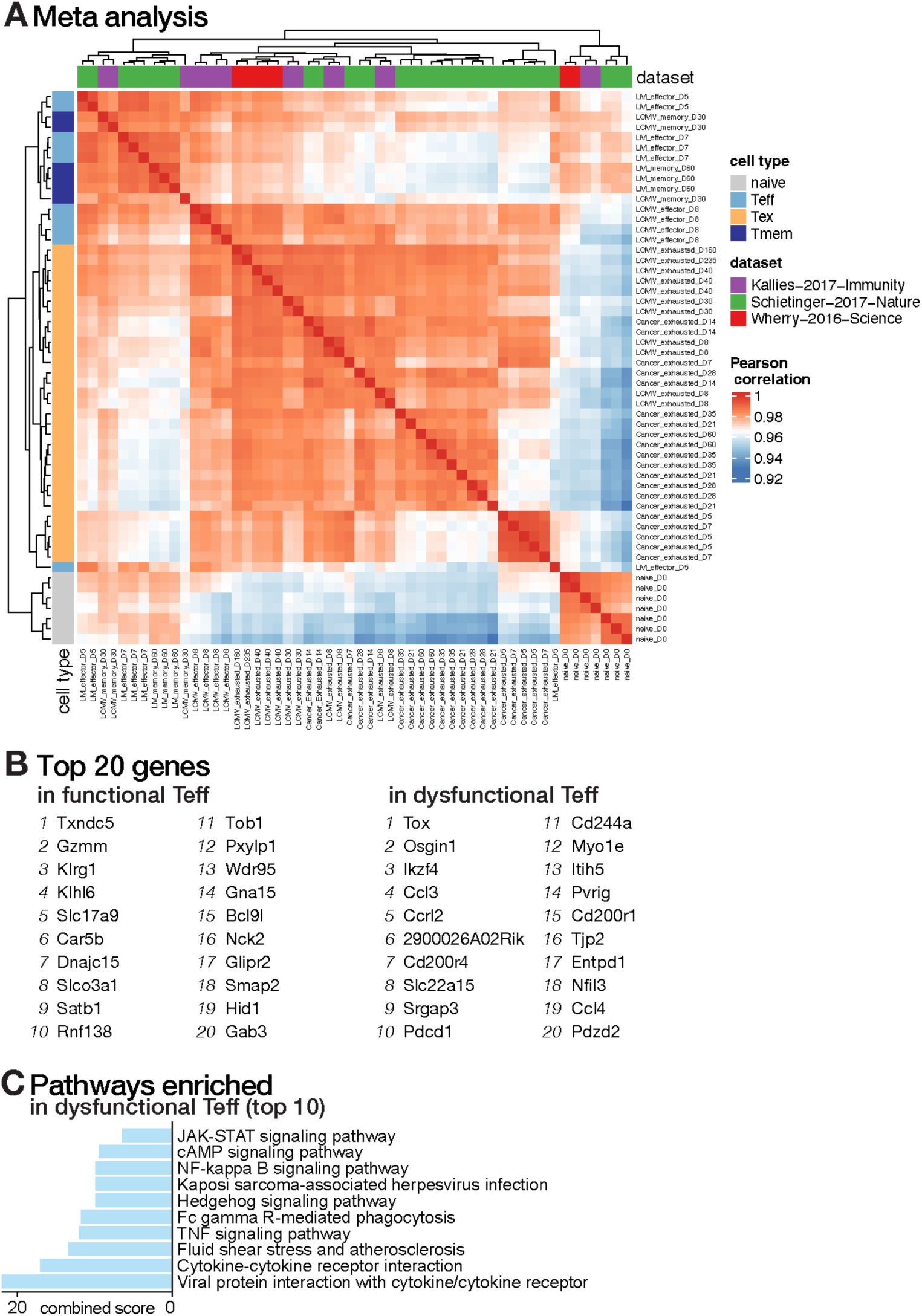
Gene expression analysis of *in vivo* generated CD8^+^ T cell differentiation states. (**a**) Full Pearson correlation-driven similarity matrix analysis of gene expression profiles of CD8^+^ T cell states (naïve, Teff – effector T cells, Tex – exhausted T cells, Tmem – memory T cells) from **Figure 1a**. Analysis was conducted on RNA-seq datasets from three independent studies characterizing gene expression profiles of antigen-specific CD8^+^ T cells from acute infection (LCMV Armstrong, *Lm*), chronic infection (LCMV CL-13), and cancer (hepatocellular carcinoma) models *in vivo*. (**b**) List of top 20 genes in functional (*left*) and dysfunction (*right*) Teff cells from rank analysis of median Wald test statistics in **Figure 1b**. for DEG between Teff and Tex populations (Teff/Tex) were calculated based on cancer and virus response datasets. (**c**) KEGG pathway analysis of the top pathways enriched in dysfunctional Teff from **Figure 1b**.

**Figure S2, related to Figure 1.**
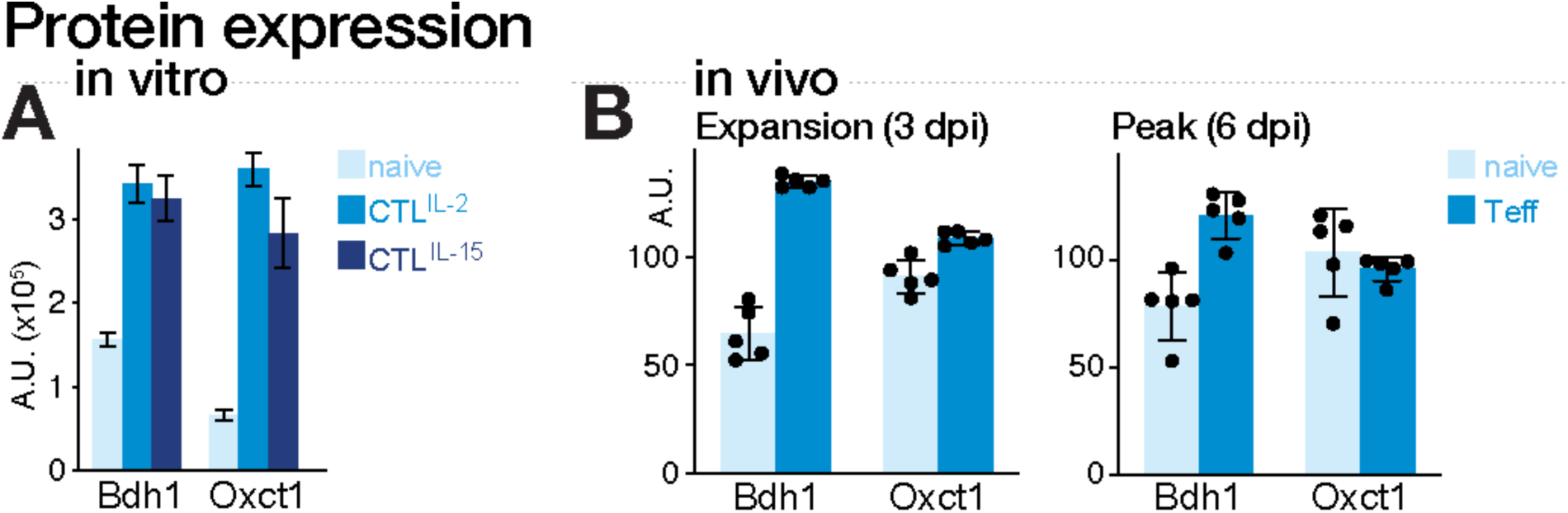
Comparison of ketolytic enzyme expression in CD8^+^ effector T cells. (**a**) Bar graphs depicting abundance of BDH1 and SCOT (Oxct1) proteins extracted from *in vitro* cultured naïve, cytotoxic (CTL^IL-2^), and memory-like (CTL^IL-15^) CD8^+^ T lymphocytes. Data were obtained from the ImmPRes proteomics resource (www.immpres.co.uk). Data represent the mean ± SEM (n=3). (**b**) Bar graphs depicting abundance of Bdh1 and SCOT/Oxct1 proteins extracted from OT-I CD8^+^ cells or bystander naïve CD8^+^ T cells isolated from *Lm-*OVA infected mice at 3 (T cell expansion phase) or 6 (peak of T cell expansion) days post infection (dpi). Data represent the mean ± SEM (n=5). A.U., arbitrary unit.

**Figure S3, related to Figure 1.**
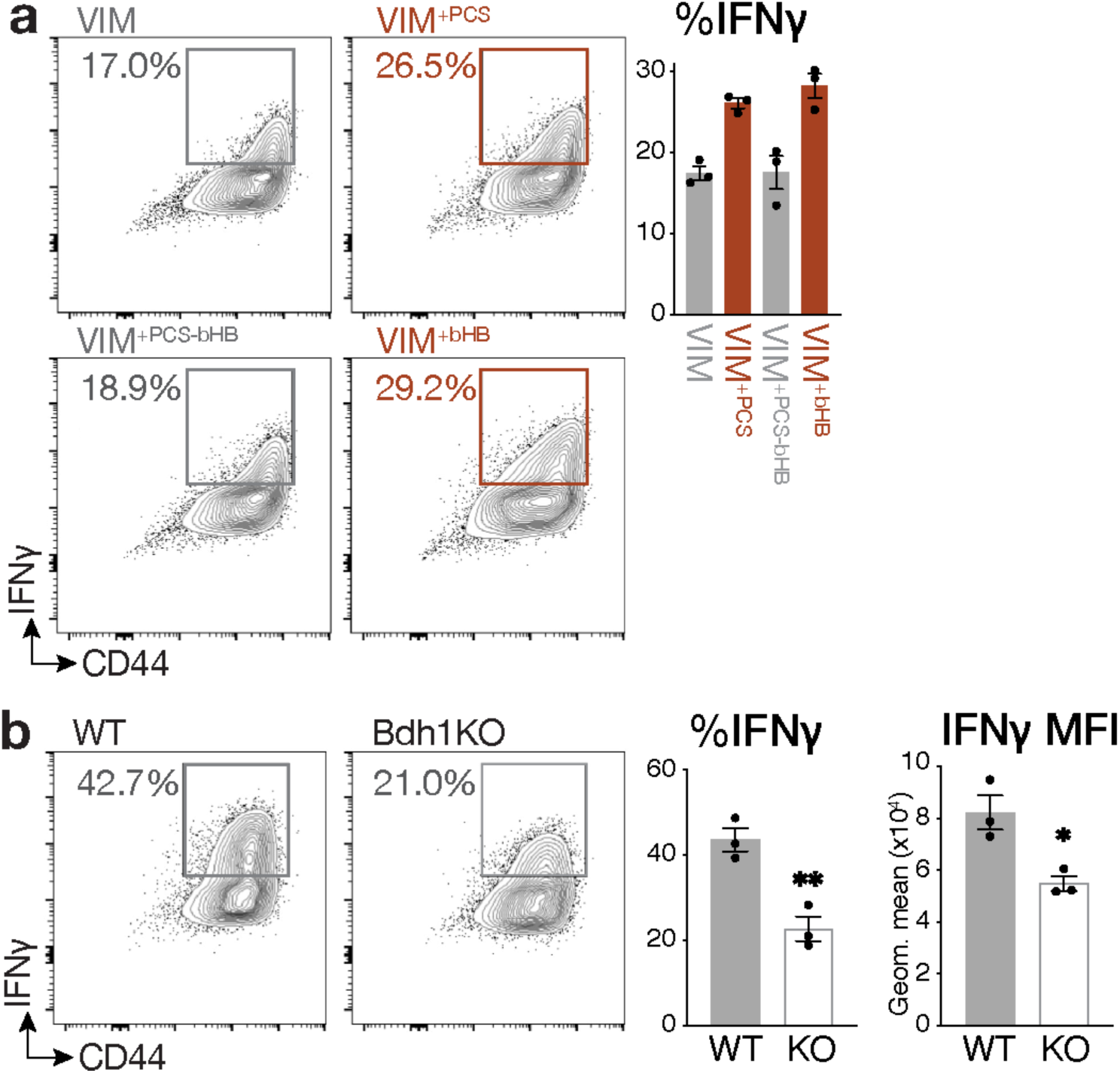
Impact of βOHB and Bdh1-deficiency on IFN-γ production in CD8^+^ Teff cells. (**a**) IFN-γ production by CD8^+^ T cells cultured in VIM, VIM containing physiologic carbon sources (PCS), VIM containing PCS minus 5 mM βOHB, and VIM with 5 mM βOHB. *Left,* Representative flow cytometry plots for CD44 versus IFN-γ expression by CD8^+^ T cells 3 days post activation. *Right,* Bar graph showing the percentage of IFN-γ^+^ T cells in each culture. Data represent the mean ± SEM (n=3). (**b**) Intracellular IFN-γ production by control (wild type, WT) and *Bdh1*-deficient (Bdh1KO) CD8^+^ T cells activated for 3 days in IMDM. *Left,* Representative flow cytometry plots for CD44 versus IFN-γ expression by WT and Bdh1KO CD8^+^ T cells 3 days post activation. *Right,* Bar graph showing the percentage of IFN-γ^+^ T cells and the geometric mean fluorescence intensity (MFI) for IFN-γ^+^ T cells. Data represent the mean ± SEM (n=3). *, *p* < 0.05; **, *p* < 0.01.

**Figure S4, related to Figure 1.**
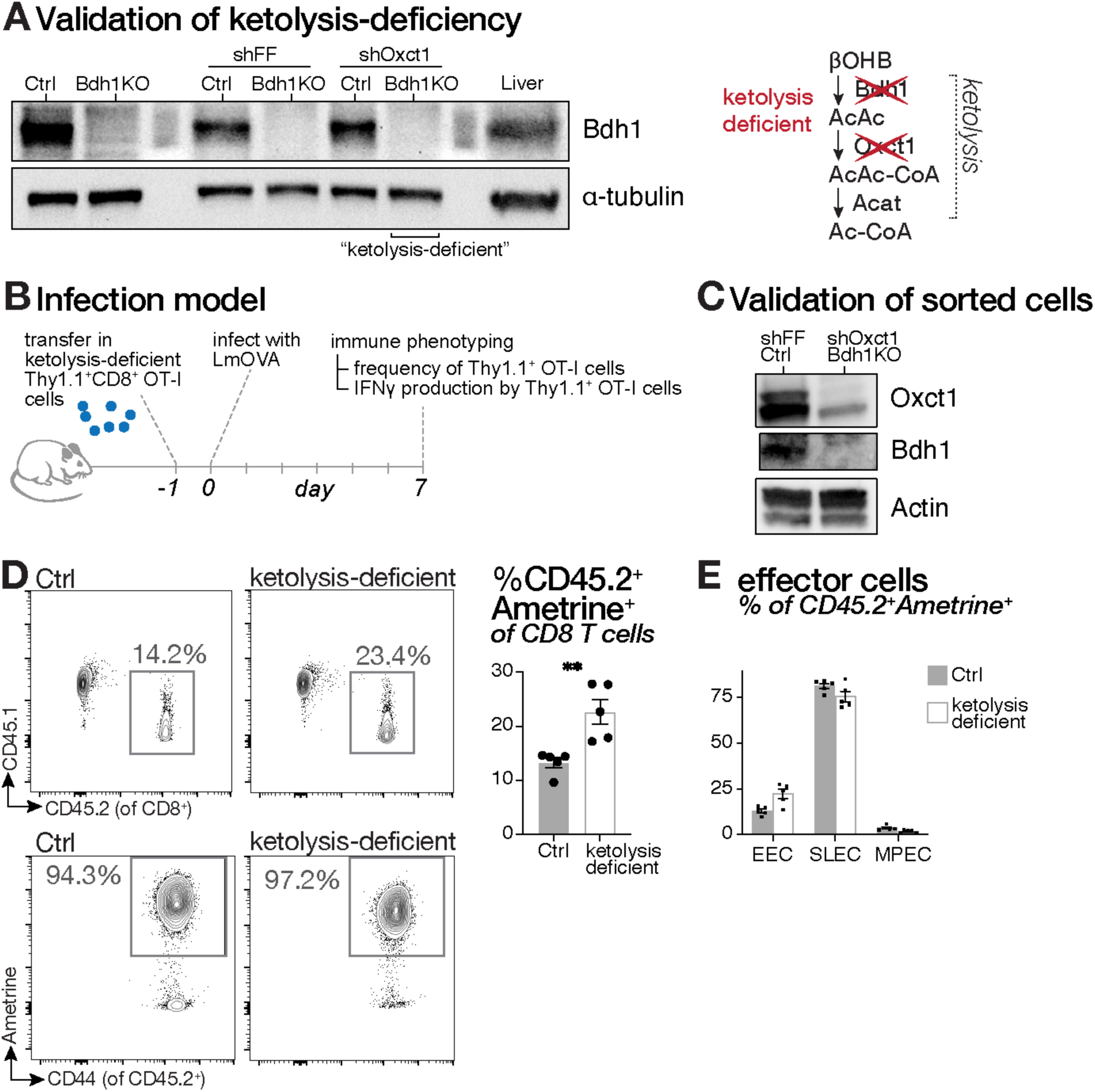
Validation of ketolysis-deficient CD8^+^ effector T cells and their *in vivo* response to *Lm*-OVA. (**a**) Validation of ketolysis-deficient T cells. *Left*, immunoblot of BDH1 protein expression in activated control (Ctrl) or Bdh1KO (*Cd4-Cre*;*Bdh1^fl/fl^*) T cells transduced with control (*shFF*, firefly luciferase) or SCOT-targeting (*shOxct1*) retroviral vectors. *Right*, schematic depicting intervention strategy for generation of ketolysis-deficient T cells. (**b**) Schematic depicting the experimental setup of the adoptive OT-I T cell transfer model coupled to *Lm-*OVA infection used for immune phenotyping at 7 dpi. (**c**) Validation of *Oxct1*-silencing in ketolysis-deficient T cells. Immunoblot of SCOT and BDH1 expression in CD8^+^ T cells from control T cells expressing shRNA control vector (*shFF*;Ctrl) versus Bdh1KO cells transduced with SCOT-targeting (*shOxct1*) retroviral vector (*shOxct1*;Bdh1KO). Actin levels are shown as a loading control. (**d**) Response of splenic OT-I CD8^+^ cells to *Lm*-OVA 7 dpi as described in **(b)**. *Left*, Representative flow cytometry plots for CD45.1 (host) and CD45.2 (donor) expression by CD8^+^ cells, and detection of Ametrine signal within the CD8^+^CD45.2^+^T cell population for the identification of adoptively transferred OT-I CD8^+^ T cells (CD8^+^CD45.2^+^Ametrine^+^). *Right*, bar graph illustrating frequency of CD45.2^+^Ametrine^+^ OT-I T cells as a percentage of total CD8^+^ T cells. Data represent the mean ± SEM (n=5). (**e**) Distribution of OT-I^+^ T cells into various differentiation states at 7 dpi with *Lm*-OVA (mean ± SEM, n=5). T cell effector states are defined as: early effector cells (EEC), KLRG1^-^CD127^-^; short lived effector cells (SLEC), KLRG1^+^CD127^lo^; early memory precursor cells (MPEC), KLRG1^-^CD127^hi^. **, *p* < 0.01.

**Figure S5, related to Figure 1.**
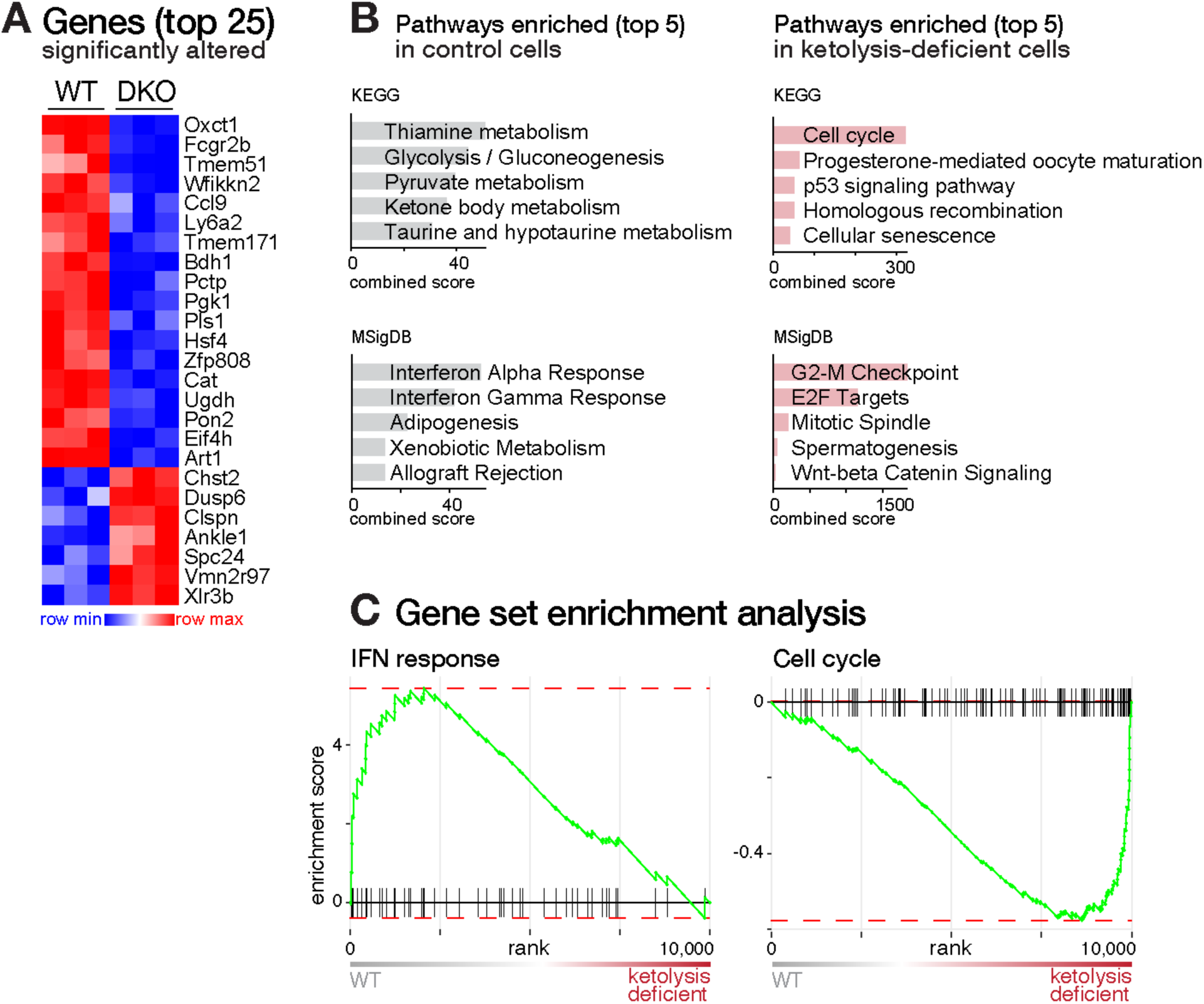
Transcriptional analysis of *Lm*-OVA-specific ketolysis-deficient CD8^+^ Teff cells. (**a**) Heatmap depicting the top 25 differentially expressed gene (DEG) transcripts (*p<0.05*) altered in control (WT) and ketolysis-deficient (DKO) OT-I CD8^+^ cells isolated from *Lm*-OVA-infected mice 7 dpi (n=3/group) (from **Figure S4**). (**b**) Bar graphs representing top five pathways from KEGG and MSigDB databases enriched in control (WT) versus ketolysis-deficient (DKO) OT-I CD8^+^ cells isolated from *Lm*-OVA-infected mice 7 dpi (n=3/group). (**c**) Gene set enrichment analysis (GSEA) of IFN response (MSigDB) and cell cycle (KEGG) genes comparing control (WT) with ketolysis-deficient OT-I T cells responding to *Lm*-OVA (7 dpi, n=3 biological replicates/group).

**Figure S6, related to Figure 2.**
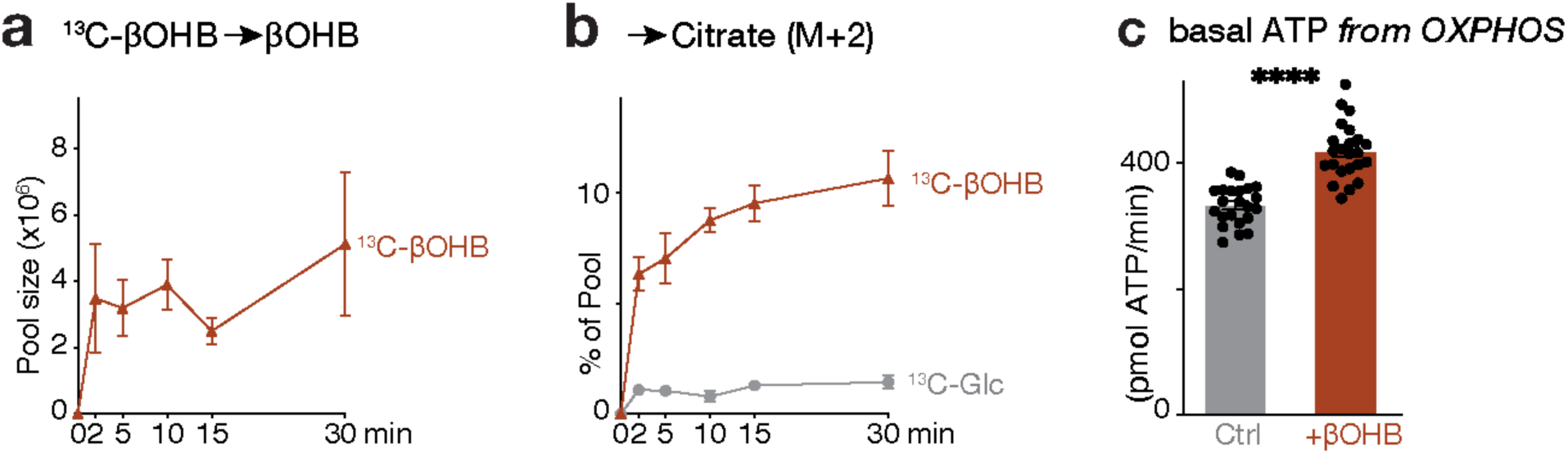
Metabolic characterization of βOHB utilization in *in vitro* activated CD8^+^ effector T cells. (**a**) Time course of βOHB pool sizes in activated CD8^+^ T cells cultured with [U-^13^C_4_]βOHB. (**b**) Relative contribution of [U-^13^C_4_]βOHB or [U-^13^C_6_]glucose to the M+2 pool of citrate in activated CD8^+^ T cells over time (mean ± SEM, n=3). (**c**) Bioenergetic profile of *in vitro* activated CD8^+^ T cells cultured in the presence of 2mM βOHB. Bar graph represents basal ATP production rate generated from OXPHOS. Data represents mean±SEM, n=22-23/group ****, p < 0.0001

**Figure S7, related to Figure 2.**
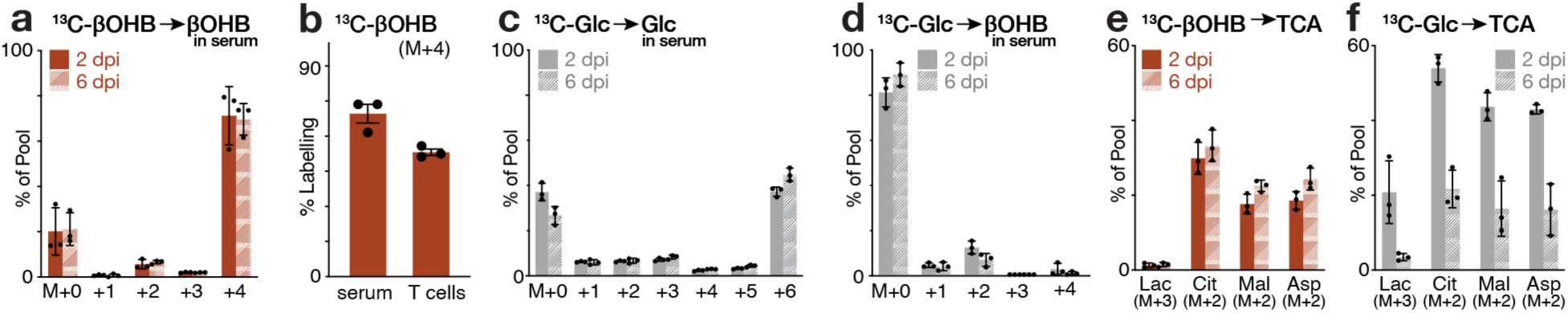
Metabolic fate of [U-^13^C_4_]βOHB and [U-^13^C_6_]glucose during *in vivo* infusions during *Lm*-OVA infection. Mice were adoptively transferred with Thy1.1^+^OT-I T cells and infected with *Lm*-OVA (see **Fig. 1G**). At 2 or 6 dpi the mice were infused with either [U-^13^C_4_]βOHB or [U-^13^C_6_]glucose for 2 h and serum and T cells harvested for analysis. (**a**) Bar graph depicting MIDs of circulating βOHB after [U-^13^C_4_]βOHB infusions at 2 and 6 dpi; M+0 represents the endogenous unlabeled βOHB pool and M+4 represents fully-labeled infused [U-^13^C_4_]βOHB. Data represent mean ± SEM for biological replicates (n=3). (**b**) Relative enrichment of [U-^13^C_4_]βOHB of the total βOHB pool in serum and the intracellular βOHB in isolated Thy1.1^+^OT-I T cells at 6 dpi (mean ± SEM, n=3). (**c-d**) MIDs of serum **(c)** glucose and **(d)** βOHB following [U-^13^C_6_]glucose infusions at 2 and 6 dpi. M+0 represents unlabeled metabolites, M+6 glucose in **(c)** represents enrichment of infused [U-^13^C_6_]glucose, and M+2 and M+4 in **(d)** represents enrichment of ^13^C-glucose-labeled βOHB due to ketogenesis. Data represent mean ± SEM for biological replicates (n=3). (**e-f**) Relative contribution of ^13^C label into lactate and TCA cycle derivatives citrate, malate and aspartate following (**e**) [U- ^13^C_4_]βOHB and (**f**) [U-^13^C_6_]glucose infusions at 2 and 6 dpi. MIDs for each metabolite were normalized relative to serum enrichment of (**e**) M+4 βOHB and (**f**) M+6 glucose, respectively. For all panels, the data represent the mean ± SEM for 3 biological replicates.

**Figure S8, related to Figure 3.**
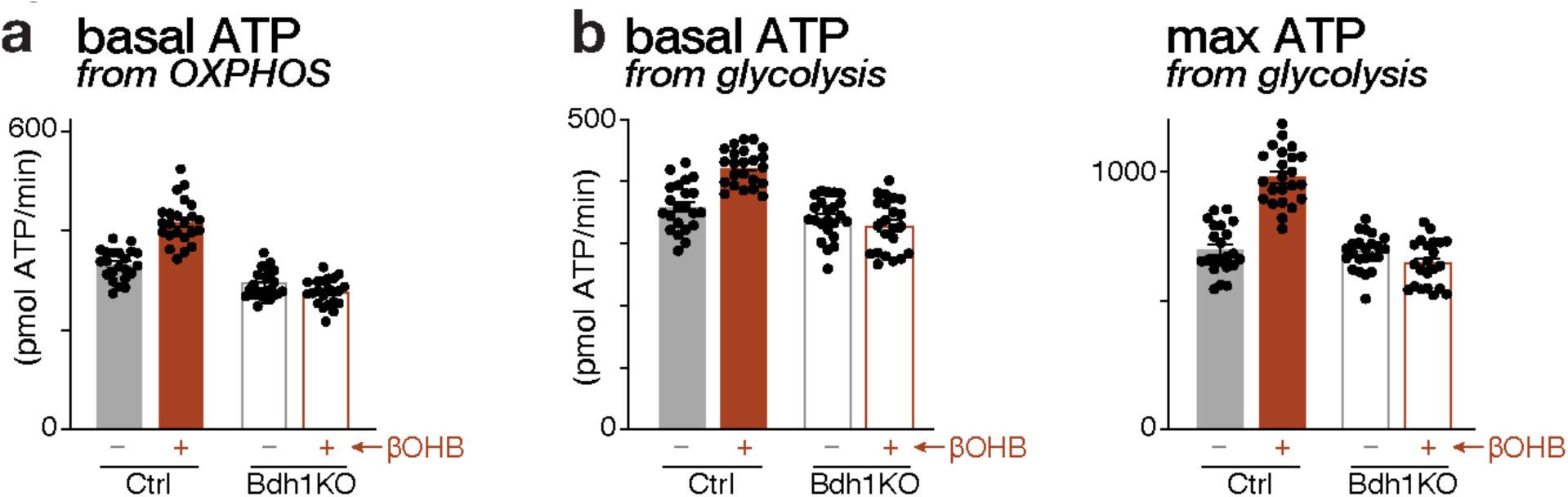
βOHB increases the bioenergetic capacity of *in vitro*-activated CD8^+^ Teff cells. Bioenergetic profile of *in vitro*-activated control (Ctrl) and Bdh1KO CD8^+^ T cells cultured in the presence (+) or absence (-) of 2mM βOHB. (**a**) Basal ATP production rates from OXPHOS. (**b**) Basal (*left*) and maximal (*right*) ATP production rates from glycolysis. Data represent the mean ± SEM, n=22-24/group.

**Figure S9, related to Figure 3.**
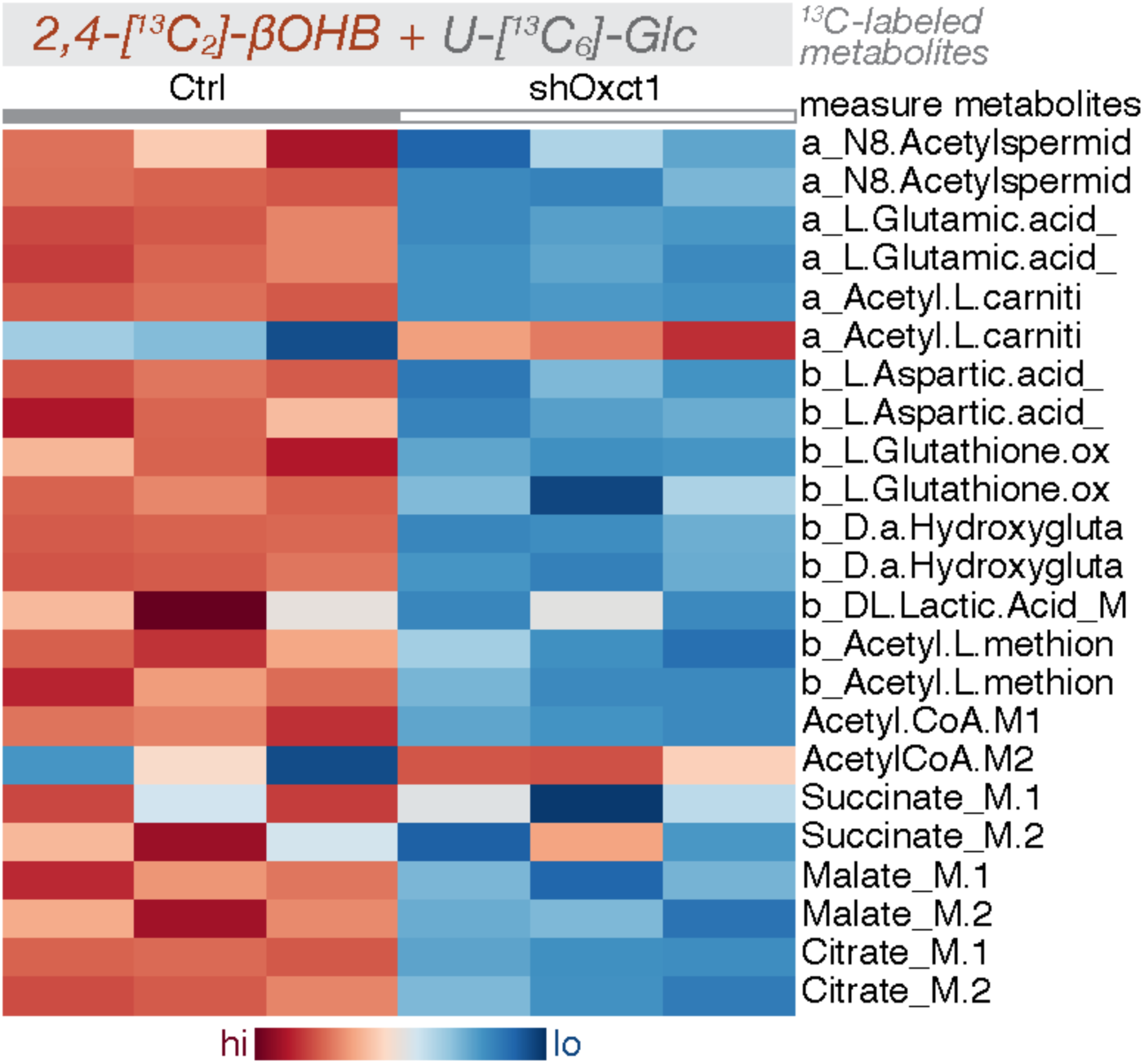
Effects of Scot1/*Oxct1*-silencing on the metabolism of [U-^13^C_6_]glucose and [2,4-^13^C_2_]βOHB in CD8^+^ Teff cells. Heatmap depicting relative enrichment of ^13^C-label in indicated metabolites extracted from *in vitro*-activated control (*shFF*) and SCOT-silenced (*shOxct1*) CD8^+^ T cells cultured in the presence of both [U-^13^C_6_]glucose and [2,4-^13^C_2_]βOHB for 2 h. For each listed metabolite, the top isotopologue (M+1) is derived from [2,4-^13^C_2_]βOHB, while the bottom isotopologue (M+2) is derived from [U-^13^C_6_]glucose (n=3/group).

**Figure S10, related to Figure 3.**
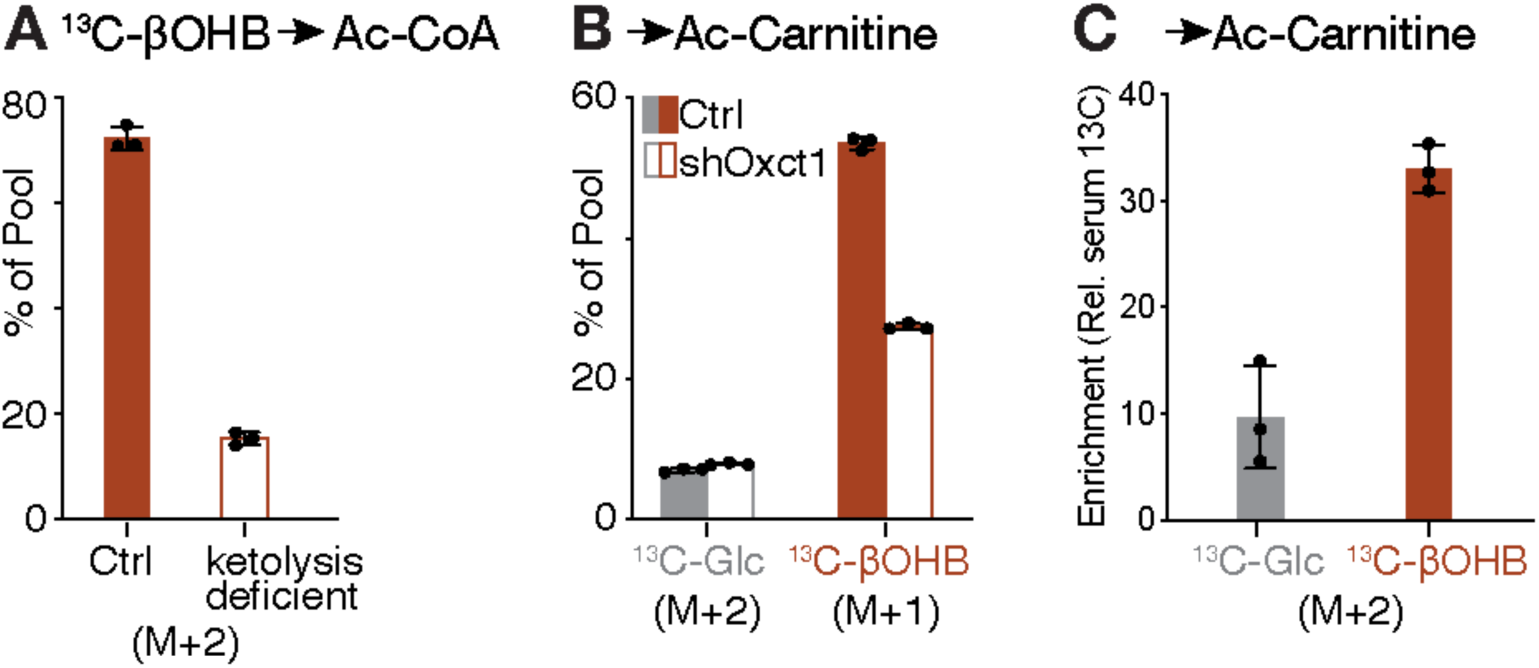
βOHB-derived carbon contributes to acetyl-CoA metabolism in activated CD8^+^ Teff cells. (**a**) Metabolic production of acetyl-CoA from [U-^13^C_4_]βOHB in activated control (Ctrl) and ketolysis-deficient (*shOxct1;*Bdh1KO) CD8^+^ T cells. Shown is fractional enrichment of the acetyl group carbon (M+2) from ^13^C-βOHB into the acetyl-CoA pool *in vitro*-activated T cells (as in Figure S4C) after 2 h of culture. Data represent the mean ± SEM (n=3 replicates/group). (**b**) Relative ^13^C contribution from glucose (M+2) or βOHB (M+1) to the acetyl-carnitine pool in activated T cells. Control (Ctrl, *shFF*) and *shOxct1*-expressing CD8^+^ T cells were cultured for 2 h with medium containing 2 mM [2,4-^13^C_2_]βOHB and 5 mM [U-^13^C_6_]glucose. Data represent the mean ± SEM (n=3/group). (**c**) Enrichment of ^13^C carbon from [U-^13^C_6_]glucose or [U-^13^C_4_]βOHB into acetyl-carnitine (M+2) in Thy1.1^+^ OT-I T cells following 2 h infusion at 6 dpi with *Lm*-OVA (as in **Figure 2j**). MIDs were normalized relative to serum enrichment of M+4 βOHB and M+6 glucose, respectively. Data represents mean ± SEM for 3 biological replicates/group.

**Figure S11, related to Figure 3.**
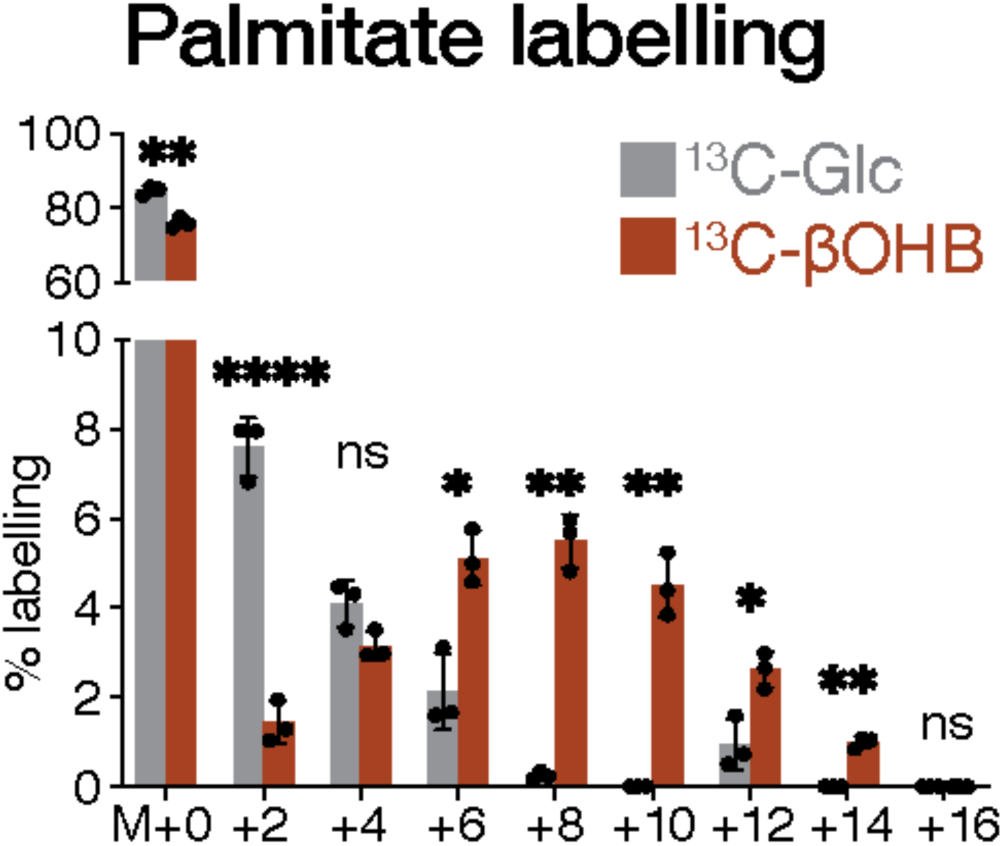
Contribution of glucose and βOHB to palmitate synthesis in CD8^+^ Teff cells. MID for [U-^13^C_6_]glucose- and [U-^13^C_4_]βOHB-derived carbon into the intracellular palmitate pool for *in vitro*-activated CD8^+^ Teff cells after 24 h of culture with each respective tracer. Data represent mean ± SEM, n=3 replicates/group. ns, not significant; *, *p* < 0.05; **, *p* < 0.001; ****, *p* < 0.00001.

**Figure S12, related to Figure 4.**
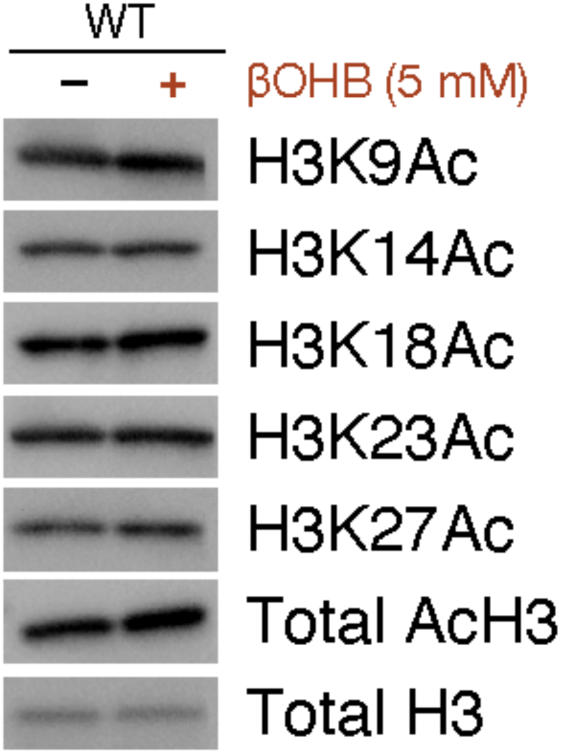
Impact of exogenous βOHB on histone H3 acetylation in CD8^+^ Teff cells. Immunoblot of total histone H3 acetylation (AcH3) and acetylation at specific histone H3 residues (K9, K14, K18, K23, and K27) in isolated histones from activated CD8^+^ T cells cultured in the presence (+) or absence (-) of 5 mM βOHB for 24 h. Total histone H3 levels are shown as a loading control.

**Figure S13, related to Figure 4.**
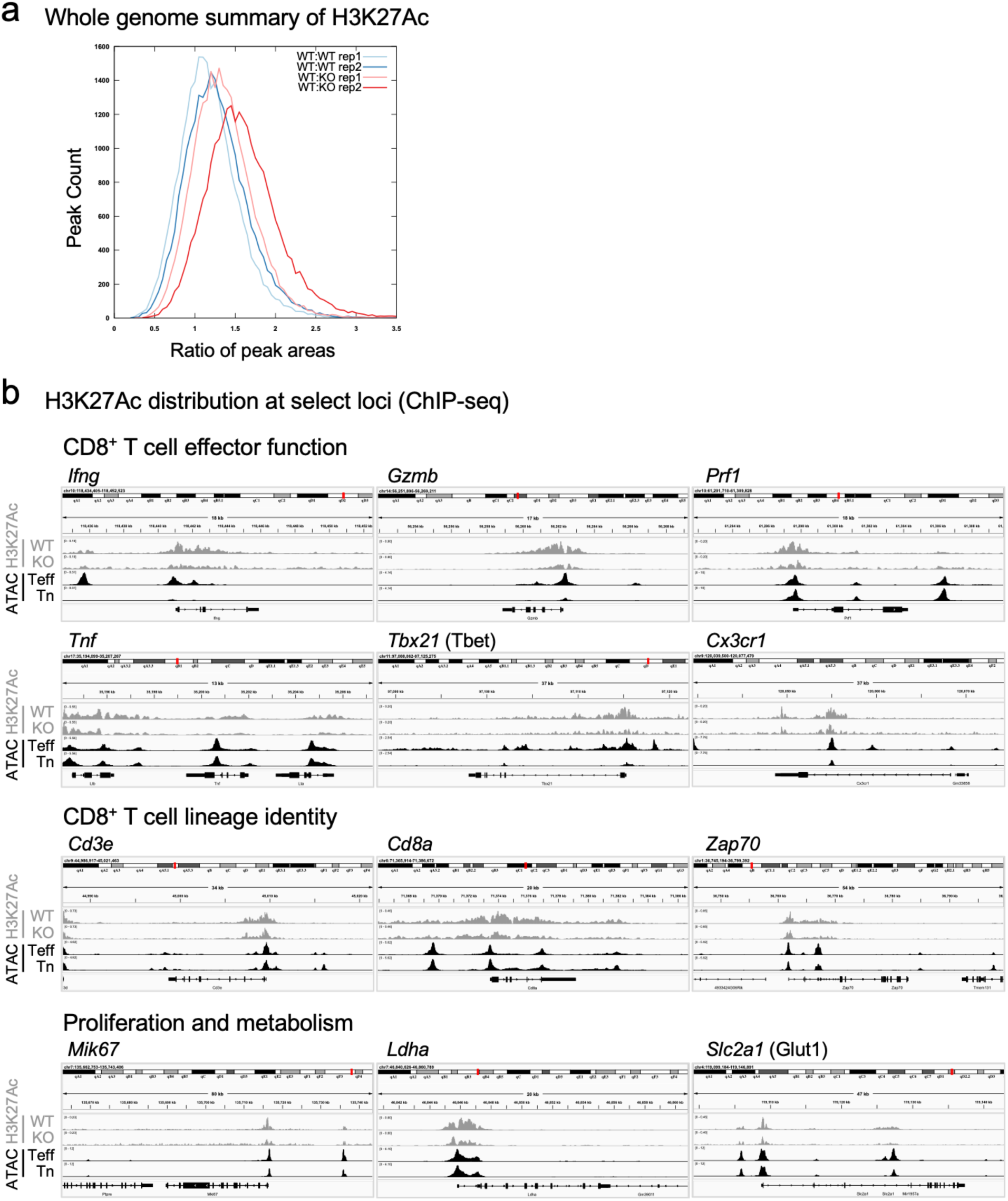
siQ-ChIP sequencing analysis of H3K27Ac enrichment at effector gene loci in CD8^+^ Teff cells. (**a**) Histogram comparing genome-wide enrichment of H3K27Ac siQ-ChIP signal (area under each peak) between control (WT) and Bdh1KO (KO) replicates. (**b**) Representative genomic tracks (full genomic loci) for H3K27Ac enrichment (grey) for genes associated with CD8^+^ effector T cell function, CD8^+^ lineage identity, proliferation, and metabolism from control (WT) and Bdh1KO CD8^+^ T cells activated for 24 h with anti-CD3 and anti-CD28 antibodies. Chromatin accessibility (ATC-seq) at corresponding genetic loci in naïve (Tn) and CD8^+^ effector (Teff) cells from *Lm*-infected mice are shown in black (data from (*27*)). Data are representative of duplicate samples.

**Table S4.**
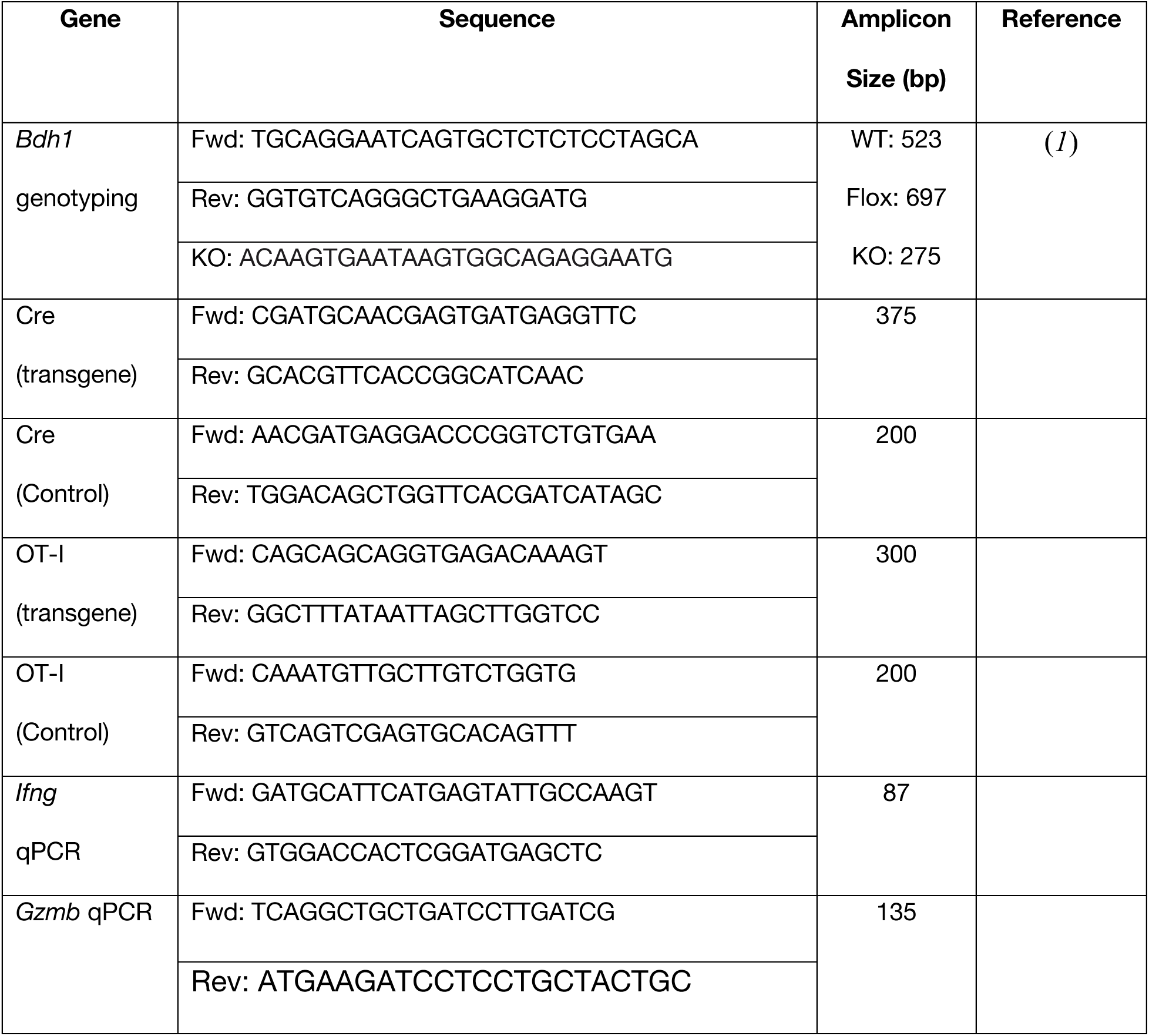
List of primers used in this study.

**Table S5.**
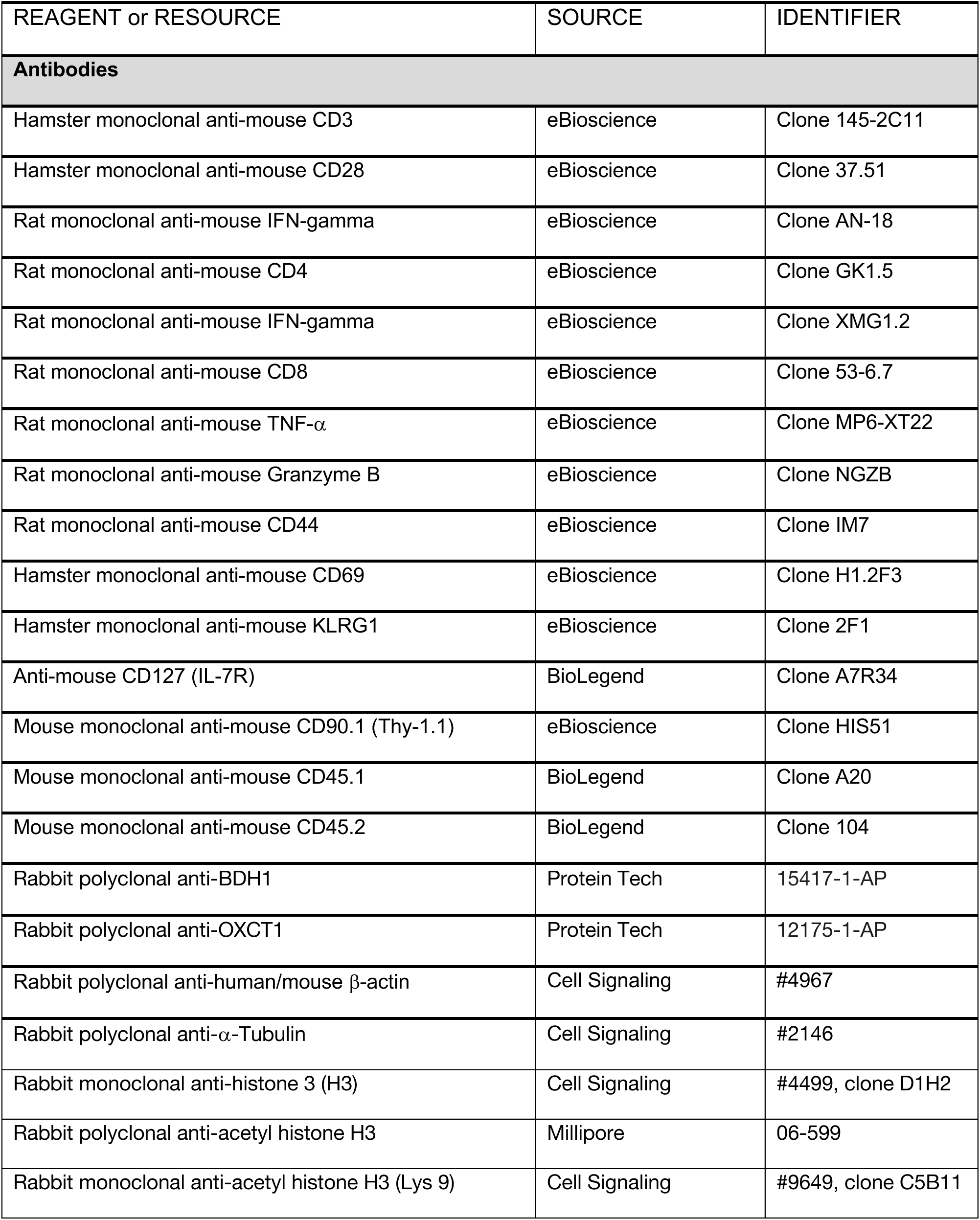
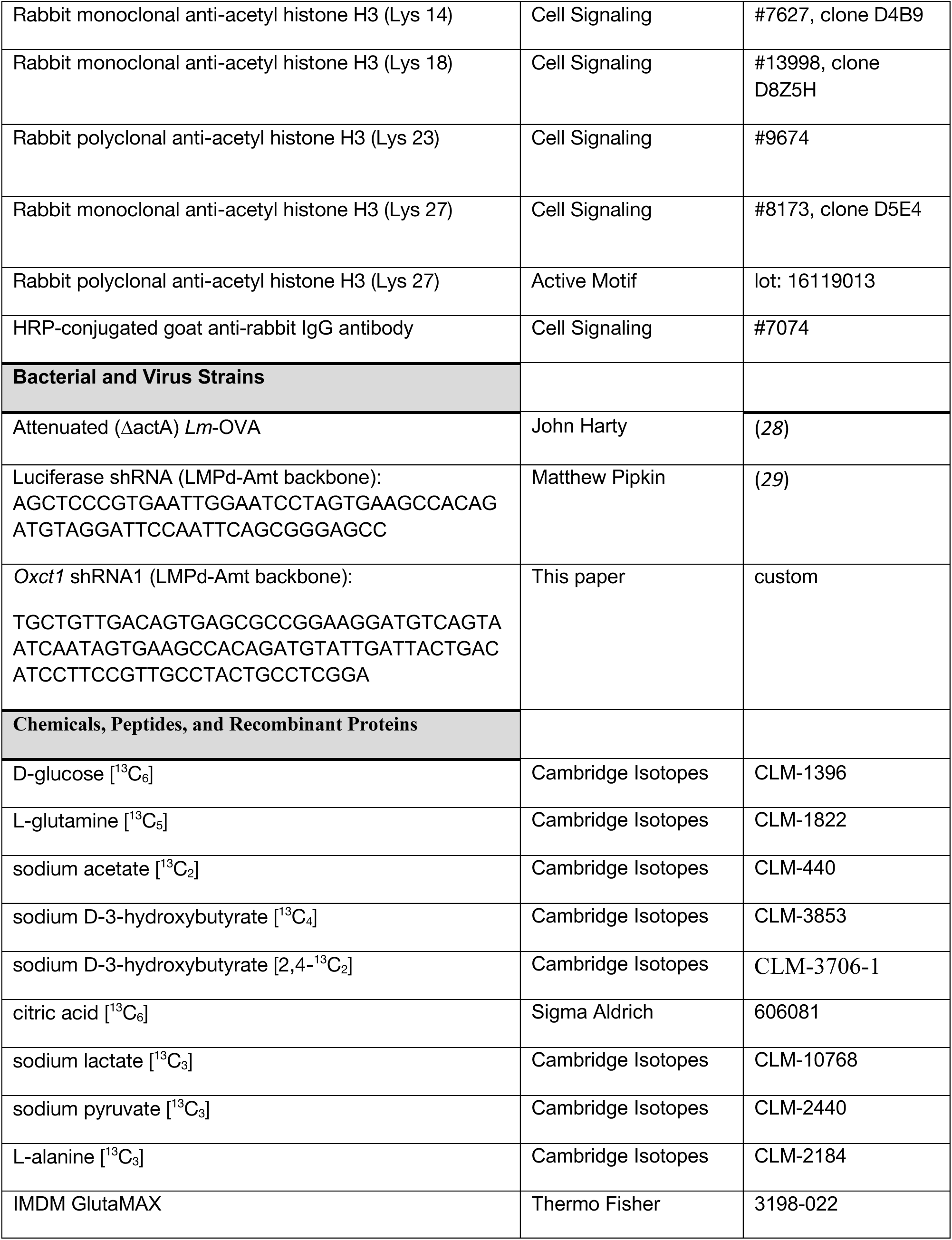
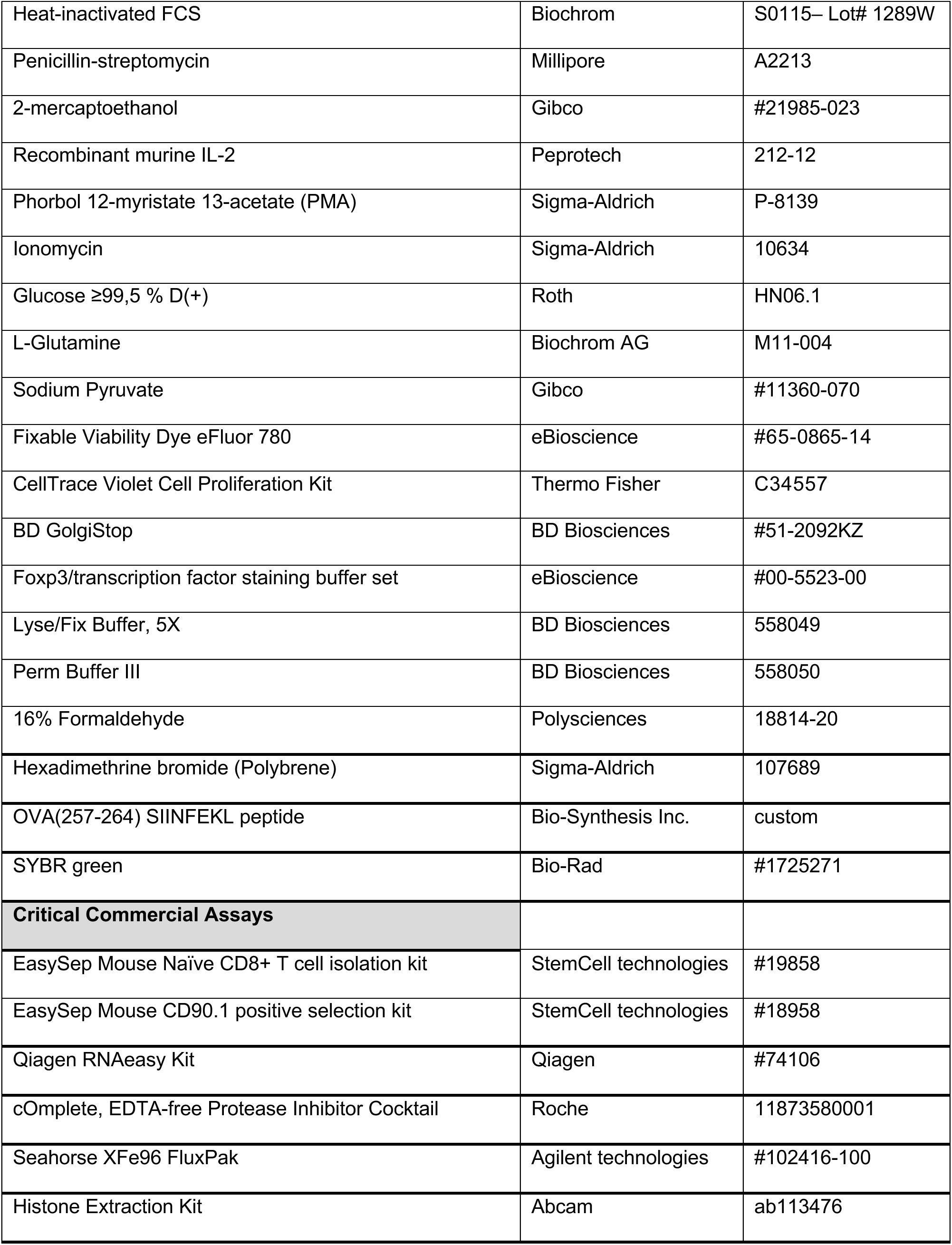
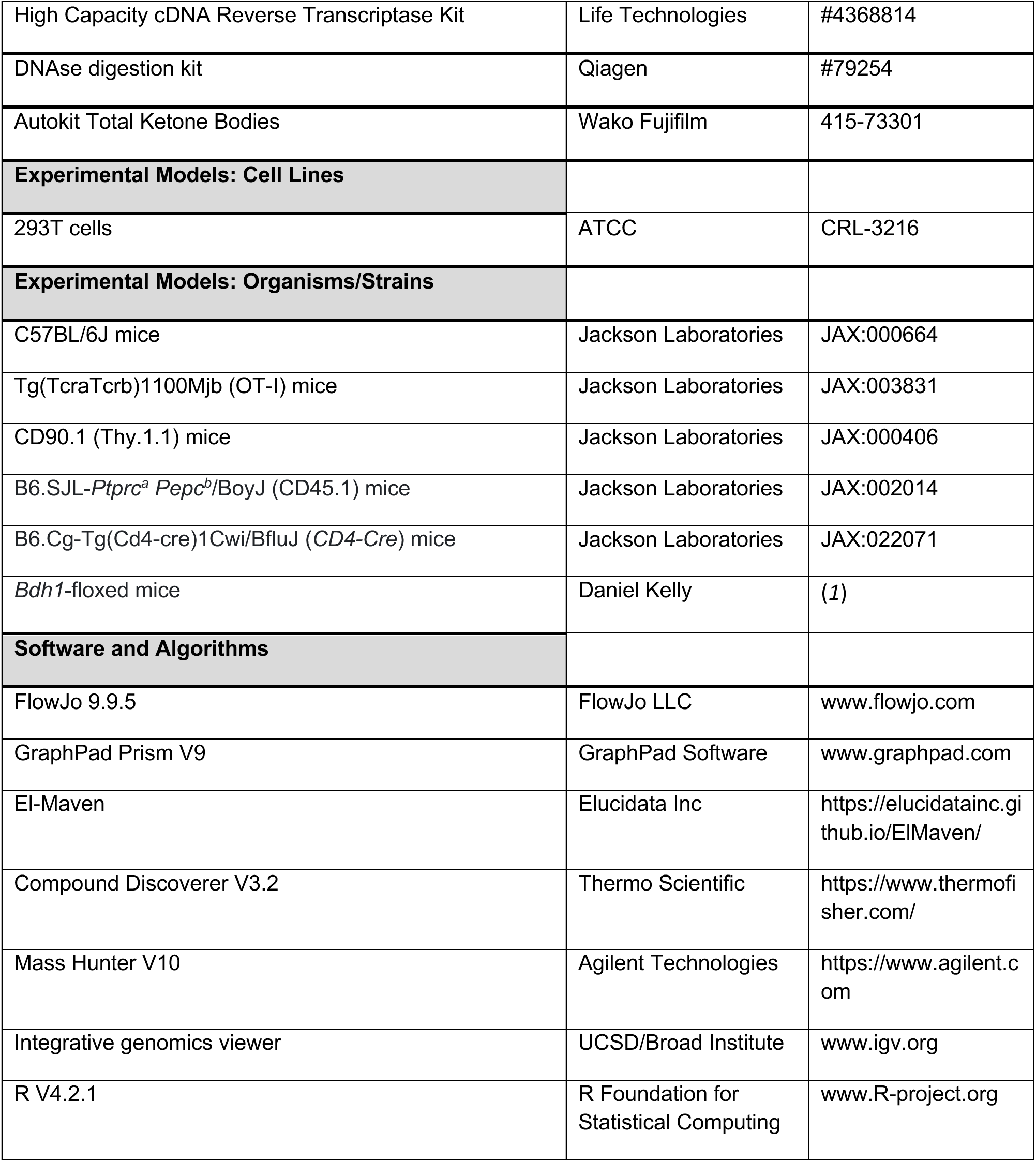
Key Resources Table.

